# Organ-level gene regulatory network models enable the identification of central transcription factors in *Solanum lycopersicum*

**DOI:** 10.1101/2025.03.26.645553

**Authors:** José D. Fernández, David Navarro-Payá, Antonio Santiago, Jonathan Canan, Sebastián Contreras-Riquelme, Ariel Cerda, Tomás C. Moyano, Lorena Melet, Nathan R. Johnson, Javier Canales, José M. Álvarez, José Tomás Matus, Elena A. Vidal

## Abstract

Tomato (*Solanum lycopersicum*) is a globally important crop, yet the gene regulatory networks (GRNs) controlling gene expression remain poorly understood. In this study, we constructed GRNs for roots, leaves, flowers, fruits, and seeds by inferring transcription factor (TF)–target interactions from over 10,000 RNA-seq libraries using the GENIE3 algorithm. We refined these networks with gene co-expression data and computational predictions of TF binding sequences in open chromatin sites. Our networks confirmed key TFs, including TOMATO AGAMOUS LIKE 1 and RIPENING INHIBITOR in fruit ripening, as well as ABF3 and ABF5 in abscisic acid response in leaves. Additionally, we identified novel candidate regulators, including AUXIN RESPONSE FACTOR 2A and ETHYLENE RESPONSE FACTOR.E2 in fruit ripening and G-BOX BINDING FACTOR 3 (*Sl*GBF3) in ABA-related and drought pathways. To further validate the GRNs, we used DNA Affinity Purification Sequencing (DAP-seq) for *Sl*GBF3, confirming the accuracy of our GRNs. This study provides a valuable resource for dissecting transcriptional regulation in tomato, with potential applications in crop improvement. The GRNs are publicly accessible through a user-friendly web platform at https://plantaeviz.tomsbiolab.com/tomviz.

**Highlight:** We developed organ-level gene regulatory networks for tomato using 10,000+ RNA-seq libraries, validated predictions and identified new regulators of fruit ripening and ABA response. These networks are available at https://plantaeviz.tomsbiolab.com/tomviz.

## Introduction

Tomato *(Solanum lycopersicum L.)* is one of the most widely cultivated and consumed crops globally. It also serves as a key model organism for research on fleshy fruit development, ripening, and plant defense responses (Kimura and Sinha, 2008; Gascuel et al., 2017). Despite its significance, the regulatory networks controlling tomato responses to internal and external signals remain largely unexplored.

Various approaches exist to help in the identification of TF-target interactions and generate gene regulatory networks. Chromatin immunoprecipitation followed by sequencing (ChIP-seq) and DNA affinity purification sequencing (DAP-seq) can directly assess events of TF binding in genomic regions; however, these methods are technically challenging, particularly in non-model organisms (Park, 2009). In *Solanum lycopersicum*, only a limited number of TFs have been studied using ChIP-seq (Fujisawa *et al*., 2011; Ricardi *et al*., 2014; Du *et al*., 2017; Lü *et al*., 2018; Gao *et al*., 2019; Lira *et al*., 2020; Liu *et al*., 2020; Ding *et al*., 2022; Tu *et al*., 2022; Yang *et al*., 2022; Jiang *et al*., 2023). Similarly, just a limited number of TFs have been studied using DAPseq (López-Vidriero *et al*., 2021; Chong *et al*., 2022; Huang *et al*., 2023; Zhu *et al*., 2023). While these approaches have identified TF binding sites and regulatory targets of TFs, they are typically limited to individual TFs. Alternatively, studies addressing accessible chromatin sites to derive binding of multiple TFs at a genome-wide scale in tomato are scarce. Most ATAC-seq analyses are focused on fruit responses to abiotic stress (Maher *et al*., 2018; Reynoso *et al*., 2019; Hendelman *et al*., 2021; Kajala *et al*., 2021), while DNase-seq studies also remain limited (Qiu *et al*., 2016; Lü *et al*., 2018).

The existing biological network models in tomato have focused primarily on protein-protein interactions or gene co-expression networks (GCNs), often derived from small datasets or limited to condition-specific analyses (Ozaki *et al*., 2010; Fukushima *et al*., 2012; Gao *et al*., 2013; Koenig *et al*., 2013; Pan *et al*., 2013; Ichihashi *et al*., 2014; Arhondakis *et al*., 2016; Kim *et al*., 2017; Zouine *et al*., 2017; Xie *et al*., 2019; Bae *et al*., 2021; Bizouerne *et al*., 2021; Kusano *et al*., 2022; Wang *et al*., 2023a; Li *et al*., 2024). Some studies have integrated larger scale transcriptomic datasets from microarrays and RNA-Seq to generate GCNs. Fukushima et al., (2012) analyzed 307 microarrays from 17 experiments, encompassing diverse conditions in a single GCN that was separated into organ-specific submatrices. Similarly, Kim et al., (2017) compiled 1,473 expression samples from 12 studies of microarrays and mRNA-Seq data to generate organ-independent coexpression networks. Zouine et al., (2017) integrated 29 RNA-seq studies to generate a global GCN. However, these networks are either organ-independent or focus only on fruits. Additionally, they rely on outdated tomato genome annotations (ITAG2.4 or ITAG3.0) (Fukushima *et al*., 2012; Kim *et al*., 2017; Zouine *et al*., 2017). Moreover, co-expression network approaches infer associations based on expression similarity rather than directed regulatory interactions (Babu *et al*., 2004; Chai *et al*., 2014; Swift and Coruzzi, 2017). Gene regulatory networks (GRNs) offer a more comprehensive framework to study TF-target interactions at a genome-wide scale. Unlike GCNs, GRNs infer directed regulatory interactions, identifying key regulatory hubs that orchestrate biological processes (Doidy *et al*., 2016; Vidal *et al*., 2020; Escorcia-Rodríguez *et al*., 2023). Therefore, developing genome-wide GRN models that integrate updated genome assemblies, diverse transcriptomic datasets, and organ-level regulation is crucial to uncovering transcriptional regulatory mechanisms in tomato.

Machine learning based approaches, such as GENIE3 (GEne Network Inference with Ensemble of Trees) have been widely used to infer GRNs from large-scale transcriptomic datasets. GENIE3 applies an ensemble of regression trees to predict TF-target interactions and has demonstrated a high performance in the DREAM4 and DREAM5 Network Inference challenges (Huynh-Thu *et al*., 2010; Huynh-Thu and Geurts, 2019). This approach has been successfully applied to generate GRNs in various plant species, including *Arabidopsis thaliana*, wheat, and maize (Huang *et al*., 2018; Harrington *et al*., 2020; Tu *et al*., 2020; De Clercq *et al*., 2021; Chen *et al*., 2023; Ranjan *et al*., 2024).In this study, we integrate large-scale tomato datasets to generate, validate, and refine organ-level GRNs. Using an extensive set of transcriptomic data and the GENIE3 algorithm, we generated five reference GRNs representing different tomato organs (roots, leaves, flowers, fruits and seeds). These networks were further enhanced with TF-target interactions derived from accessible chromatin data, predicted TF binding events, and genome-wide GCNs. The resulting organ-level GRNs provide a robust foundation for exploring diverse biological contexts, from developmental processes to stress responses, offering a valuable resource for addressing unresolved questions in tomato biology. The organ-level GRNs are available at https://plantaeviz.tomsbiolab.com/tomviz.

## Materials and Methods

### Tomato gene annotation update

Gene models from tomato annotation iTAG 4.2 beta (provided by Dr. Surya Saha, SolGenomics) were merged with gene models from iTAG4.0. New gene models reported in iTAG4.2 beta were incorporated into the iTAG4.0 annotation when their genome coordinates did not overlap existing entries, resulting in the final annotation file utilized in this work (hereafter iTAG4-merge). To expand the functional annotations for tomato, ITAG 4.0c protein sequences were analyzed using eggNOG-mapper (Cantalapiedra *et al*., 2021) to predict Gene Ontology (GO) terms, functional categories, and orthologs relationships. Additionally, functional annotations for all genes in ITAG 4.0c were generated using InterProScan v5.57-90 (Jones *et al*., 2014) under default parameters and complemented with GO information for *Solanum lycopersicum* from PLAZA 5.0 (Van Bel *et al*., 2022) database. The resulting GO annotations for molecular functions and biological processes were consolidated and used to create an updated GO set.

To update the list of TFs in ITAG 4.0c, we integrated evidence from multiple sources: 1. Gene descriptions for iTAG4.0 and iTAG4.1 available at SolGenomics (description contains the term “transcription factor”) (Fernandez-Pozo *et al*., 2015; Hosmani *et al*., 2019); 2. TF catalogs for *S. lycopersicum* obtained from PlantTFDB (Jin *et al*., 2017) and ITAK (Zheng *et al*., 2016); 3. ITAG 4.1c GO annotations (description contains the terms “transcription factor”, “DNA-binding”); The output of an OrthoFinder v3.0 (Emms and Kelly, 2019) analysis of Arabidopsis TFs from PlantTFDB; and 4. A search of proteins that we annotated with an IPR code associated to eukaryote TFs (list of IPR codes obtained from InterPro and manually curated to exclude IPR codes not representing TFs) (Jones *et al*., 2014). Genes supported by at least three independent lines of evidence were selected and a manual curation step was conducted to exclude proteins associated with enzymatic activities, non-transcriptional molecular processes (e.g., DNA replication, repair, splicing, translation), transcriptional regulators other than TFs (e.g., basal transcription factors, RNA polyadenylation factors), and chromatin remodeling complex subunits. We associated Position Weight Matrices (PWMs) to the TFs by combining information obtained from CisBP v.2 (Weirauch *et al*., 2014) with PWM inference from protein sequences using the JASPAR profile inference tool “infer_profile.py” (Castro-Mondragon *et al*., 2022).

### Processing of RNA-seq data

To obtain tomato RNA-seq datasets, we queried the NCBI SRA database using ("*Solanum lycopersicum*”[Organism] AND ILLUMINA[Platform]) NOT (RIP-Seq[Strategy] OR OTHER[Strategy] OR ChIP-Seq[Source] OR METATRANSCRIPTOMIC[Source] OR Bisulfite-Seq[Strategy] OR GENOMIC[Source] OR METAGENOMIC[Source] OR DNase-Hypersensitivity[Strategy] OR WGS[Strategy] OR ncRNA-Seq[Strategy] OR WCS[Strategy] OR degradome OR miRNA-Seq[Strategy] OR small RNA[Title] OR sRNA[Title]) (Leinonen *et al*., 2011). The metadata was classified by organ of origin following the protocol in Santiago *et al*. (2024). The libraries were downloaded using SRAtools (Kans, 2010). Adapters were trimmed, and low-quality reads (reads with average quality inferior to q<30 and shorter than 20 bases) were filtered out using fastp v.0.20.0 (Chen *et al*., 2018). Reads were aligned to the SL4.0 *S. lycopersicum* genome assembly (Hosmani *et al*., 2019) using STAR v.2.7.3 (Dobin *et al*., 2013). After mapping, a total of 10,618 mRNA-seq libraries were retained. Gene counts were obtained with FeatureCounts v.2.0.0 (Liao *et al*., 2014) using the ITAG4-merge annotation. Total counts were normalized to transcripts per million (TPM). Finally, genes with ≥ 5 TPM in at least 10% of the total libraries for each organ were classified as expressed, as described in Huang et al., (2018).

### Processing of ChIP-seq data

The following query (("*Solanum lycopersicum*”[Organism] AND ILLUMINA[Platform]) AND ChIP-Seq[Source]) was used to obtain tomato TF-binding (ChiP-seq) datasets from the NCBI SRA website (Leinonen et al., 2011). ChIP-seq libraries were processed using the methods described by ENCODE (Hitz *et al*., 2023). Briefly, the libraries were downloaded from the NCBI SRA using SRAtools (Kans, 2010). Adapters were trimmed, and low-quality reads (reads with average quality q<30 and shorter than 20 bases) were filtered using Cutadapt v.4.9 (Martin, 2011). Each file was mapped with Bowtie2 v.2.54 (Langmead and Salzberg, 2012) to the SL4.0 assembly (Hosmani *et al*., 2019). Alignment files were sorted and filtered with Samtools v.1.21 (Li *et al*., 2009) and peaks were identified with MACS2 v.2.2.9.1 (Zhang *et al*., 2008). Only libraries with ≥80% mapping efficiency and over 1,000 peaks assigned to annotated genes were retained as high-quality datasets for downstream analysis.

### Processing of ATAC-seq and DNAse-seq data

The following query (("*Solanum lycopersicum*”[Organism] AND ILLUMINA[Platform]) AND ATAC-Seq[Source] AND DNAse-Seq[Source]) was used to obtain tomato open chromatin sites (OCSs) datasets from the NCBI SRA website (Leinonen *et al*., 2011). A total of 183 open chromatin experiments (DNase-seq, ATAC-seq libraries) were downloaded using SRAtools (Kans, 2010). Reads were trimmed of adapters, and low-quality reads (reads with average quality q<30 and shorter than 20 bases) were filtered using Cutadapt v.4.9 (Martin, 2011). The ATAC-seq libraries were processed following the protocol described in Reynoso et al., (2019), while the DNase-seq libraries were processed following the protocol described in Moyano et al., (2021). Briefly, reads were mapped to the SL4.0 genome assembly (Hosmani *et al*., 2019) using Bowtie2 v.2.54 (Langmead and Salzberg, 2012). The ATAC-seq alignments were sorted and filtered with Samtools v.1.21 (Li *et al*., 2009) and peaks were identified with HOMER v.4.11 (Heinz *et al*., 2010). The DNase-seq regions were mapped into DNase hypersensitive sites using HOTSPOT (Meuleman *et al*., 2020). OCS peak files were then merged by experiment and converted into FASTA sequences with BedTools v.2.31.1 (Quinlan and Hall, 2010).

### Determination of TF binding sites using FIMO

TF-target datasets were generated by mapping TF DNA binding preferences, represented as Position Weight Matrices (PWMs) to genomic regions of the *Solanum lycopersicum* SL4.0 genome assembly using the Find Individual Motif Occurrences (FIMO) search tool, with default parameters (p-value < 1×10e^-4^) (Grant *et al*., 2011). As query sequences, we used all promoter sequences represented as two kilobases upstream of the transcription start site (TSS) of each gene, or organ-level OCS sequences. For OCS sequences, the results were assigned to genes using the BedTools command ClosestGene (Quinlan and Hall, 2010).

### GENIE3 inference of regulatory interactions

Raw gene counts for each organ, as well as the updated list of TFs, were given as input to the GENIE3 algorithm (Huynh-Thu *et al*., 2010). The GENIE3 tool was run with standard parameters, using 2,000 decision trees and a seed of 122 for reproducibility. The output scores were used to create subnetworks based on the top 1%, 2%, 5%, 8%, and 10% scores of TF-target pairs. These thresholds were used in previously published GENIE3 networks (Cuesta-Astroz et al., 2021; J.Huang et al., 2018; Olivares-Yañez et al., 2021).

### Co-expression network generation

To generate co-expression networks for tomato organs, we used the pipeline described in Orduña, et al. (2023). Briefly, the gene count tables from the transcriptome of each organ were used to calculate the Pearson correlation values for each pair of genes. The results were ordered by gene rank in descending order and computed in a highest reciprocal ranking (HRR) matrix per organ following the formula: HRR (A,B) = max (rank (A,B), rank (B,A)). Finally, to avoid noise and low confidence pairings, the top 1% of the highest frequency HRR per gene were chosen to make the tomato organ-specific GCNs.

### Evaluation of network performance

To evaluate the performance of the predicted GRNs in capturing experimentally validated regulatory interactions, we followed the protocol in Contreras-López et al., (2022) to compute the area under the receiver operating characteristic (AUROC) and precision-recall (AUPR) curves. Briefly, these analyses were performed using organ-level GRNs and tested against validated regulatory interactions derived from ChIP-seq datasets. Gene interactions were filtered to retain only regulatory genes present in both the GENIE3-inferred and ChIP-seq networks, with edges assigned as binary labels indicating the validation status. True and false positive rates were calculated using the *precrec* v.0.14.4 package in R. To assess statistical significance, the AUROC and AUPR values were compared against 1,000 randomized networks generated by shuffling edge weights. Percentile-based confidence intervals (2.5–97.5%) were used to benchmark the performance of the GENIE3 network, and significance was determined via a permutation test.

### Visualization of GRNs and network analysis

Network visualizations were generated using Cytoscape v.3.10.1 (Shannon *et al*., 2003) and network topology analyses were conducted using the Cytoscape NetworkAnalyzer tool. The R package Influential v. 2.2.9 (Salavaty *et al*., 2020) identified the TF hubs with the highest integrated value of influence (IVI).

### Gene Set Enrichment Analysis (GSEA)

Gene Set Enrichment Analysis (GSEA) was conducted to identify overrepresented biological process GO terms using a hypergeometric test with Benjamini and Hochberg false discovery rate (FDR) correction (threshold < 0.05). The analysis was performed using the BinGO v.3.0.5 tool (Maere *et al*., 2005) within Cytoscape, with input from the updated tomato 4.1c GO terms catalog. The REVIGO v. 1.8.1 (Supek *et al*., 2011) web application was used to improve and narrow down the grouping of GO terms, and GO terms within levels 5–7 were selected to focus on more specific terms.

### Generation of DAP-seq libraries

Genomic DNA (gDNA) was extracted from fully expanded mature leaves of 3-month-old *Solanum lycopersicum* cv. Moneymaker plants using the Wizard Genomic DNA Purification Kit (Promega), according to the manufacturer’s instructions. gDNA was fragmented to an average size of ∼200 bp using a M220 sonicator (Covaris, Woburn, MA, USA). The fragmented gDNA underwent end repair, A-tailing and ligation of Illumina adapters.

The full-length coding sequence of *SlGBF3* (*Solyc03g120460*) was amplified from cDNA obtained from *S. lycopersicum* cv. Moneymaker leaves using primers SlGBF3_Fw (5’-CACCATGGGAAATAGTGAGGATGGGAAATCATGTAAGC-3’) and SlGBF3_Rv (5’-TCACCCAGCTGCTACTGCATCA-3’). PCR products were cloned into the pFN19K_HaloTag-T7-SP6 Flexi expression vector (Promega, Madison, WI, USA), with the Halo-Tag at the N-terminal. Expression of the Halo-tagged fusion protein was performed using the TNT SP6 Coupled Wheat Germ Extract System (Promega). HaloTag-ligand conjugated magnetic beads (Promega) were used to pull-down the Halo-tagged TF. Pulled-down TFs were exposed to adapter-ligated gDNA libraries. Bound DNA was eluted and sequencing libraries were generated by PCR amplification with Illumina TruSeq Universal and Index primers. An empty expression vector was used as a negative control to account for non-specific DNA binding (input library). Libraries were sequenced on an Illumina NovaSeq 6000 platform (150 pb PE, approximately 22 million reads per library). Two replicates were used for all experiments.

### Processing of DAP-seq data

Raw sequence data were processed using the pipeline described by *Hutin et al., (2023)*. Briefly, adapter sequences were trimmed, and low-quality reads (reads with average quality q<30 and shorter than 20 bases) were filtered using Cutadapt v.4.9 (Martin, 2011). Filtered reads were mapped to the *Solanum lycopersicum* genome assembly SL4.0 (Hosmani *et al*., 2019) using Bowtie2 v2.5.4 (Langmead and Salzberg, 2012). Alignment files were sorted, filtered, and deduplicated using Samtools v1.21 (Li *et al*., 2009). Peaks were identified using MACS2 v2.2.9.1 (Zhang *et al*., 2008) with the parameters -f BAMPE -q 0.00001 --call-summits. Final peak-associated regions were annotated using BedTools closestGene function (Quinlan and Hall, 2010). Binding motifs were identified using the MEME-ChIP tool v5.1.1 (Machanick and Bailey, 2011).

## Results

### Refining tomato gene annotation and transcription factor catalog for GRN construction

As a first step to generate organ-level GRNs for tomato, we obtained a revised gene annotation for genome SL4.0 from SolGenomics (iTAG4.2 beta). To include all the genes, present in iTAG4.0 and include new genes, we merged the iTAG4.0 and iTAG4.2 beta versions, into the iTAG4-merge version. This version contains a total of 37,467 genes (Supplementary Datasets 1,2,3). Considering that only 15.53% of tomato genes had a functional annotation in ITAG4.0, we generated an updated annotation for protein coding genes by integrating different sources of evidence. These included obtaining GO terms assigned by EggNOG-mapper (Cantalapiedra *et al*., 2021) and InterProScan (Jones *et al*., 2014) and gathering gene functional annotations compiled in PLAZA 5.0 for tomato (Van Bel *et al*., 2022). The GO annotation in ITAG 4.0c covers 70.47% (26,403 out of 37,467) of protein-coding genes, each associated with at least one GO term. This dataset includes 9,379 unique GO terms, providing biological function annotations for 21,630 genes and molecular function annotations for 23,809 genes, for a total of 982,370 annotations (Supplementary Table S1 and S2).

To generate an updated list of TFs, we retrieved TF catalogs from various repositories, we conducted orthologous TF analyses using the set of *Arabidopsis thaliana* TFs and we obtained information from the GO annotations. Only TFs supported by at least three lines of evidence were selected, resulting in a set of 1,840 TFs for further analyses (Supplementary Table S3). DNA-binding preferences were determined by retrieving position weight matrices (PWMs) from CisBP v.2 (Weirauch *et al*., 2014) or assigning PWMs from Arabidopsis and Maize orthologs in JASPAR (Castro-Mondragon *et al*., 2022), yielding 846 TFs with assigned PWMs (Supplementary Table S3). This updated annotation and TF list served as the foundation for constructing the GRNs in the subsequent steps.

### Tomato genes exhibit widespread expression, yet their expression levels vary across different organs

We started our analysis by compiling a comprehensive dataset of publicly available tomato RNA-Seq libraries (Supplementary Table S4). The transcriptomic libraries were categorized into five main organs: roots (1,840 libraries from 124 studies), leaves (3,778 libraries from 279 studies), flowers (568 libraries from 55 studies), fruits (4,149 libraries from 147 studies), and seeds (270 libraries from 13 studies) for a total of 10,605 libraries that surpassed quality filters (Supplementary Fig. S1A). These libraries encompass a diverse range of tomato genotypes and growth conditions. Additionally, they represent multiple experimental contexts, including abiotic stress (3,379 runs), biotic interactions (2,085 runs), developmental studies (2,872 runs) and genetic modification studies (2,035 runs) (Supplementary Fig. S1B). This comprehensive dataset provides a robust foundation for constructing reference GRNs to address multiple research questions.

Reads were mapped to gene models using the ITAG4-merge annotation, and genes with expression levels above 5 TPM in more than 10% of the total libraries for a given organ were considered as expressed in that organ. We identified 26,910 genes (71.82%) as expressed in at least one organ, from which 19,361 (71.95%) genes are expressed in all organs, while a minor fraction (15.9/% of expressed genes) exhibit organ-level expression (Fig. 1A, Supplementary Table S5). Similar expression patterns have been reported in maize, flaxseed and wheat (Huang et al., 2018; Qi et al., 2023; Ramírez-González et al., 2018). Examples of specific genes include *Solyc06g051770* and *Solyc10g047720* in seeds, whose Arabidopsis homologs, *Oleosin 1* and *2 (OLEO1-2)*, are involved in seed oil body formation (Siloto *et al*., 2006). In roots, we identified *SULTR1;1* (*Solyc10g047170*), which encodes a sulfate transporter associated with external sulfate uptake (Takahashi *et al*., 2000). In flowers, *Tapetum Determinant 1-like (TPD1-like)* genes such as *Solyc11g005500*, *Solyc12g009850*, *Solyc05g010190* and *Solyc04g071640* were specifically expressed, consistent with their role in tapetal cell development and gametogenesis (Ezura *et al*., 2017). In leaves, we found *Longifolia* 1 (*SlLNG1*, *Solyc02g089030)*, whose Arabidopsis homolog *LNG1 (AT5G15580*) regulates leaf morphology by promoting longitudinal cell expansion (Lee *et al*., 2006) (Fig. 1A).

**Fig. 1.**
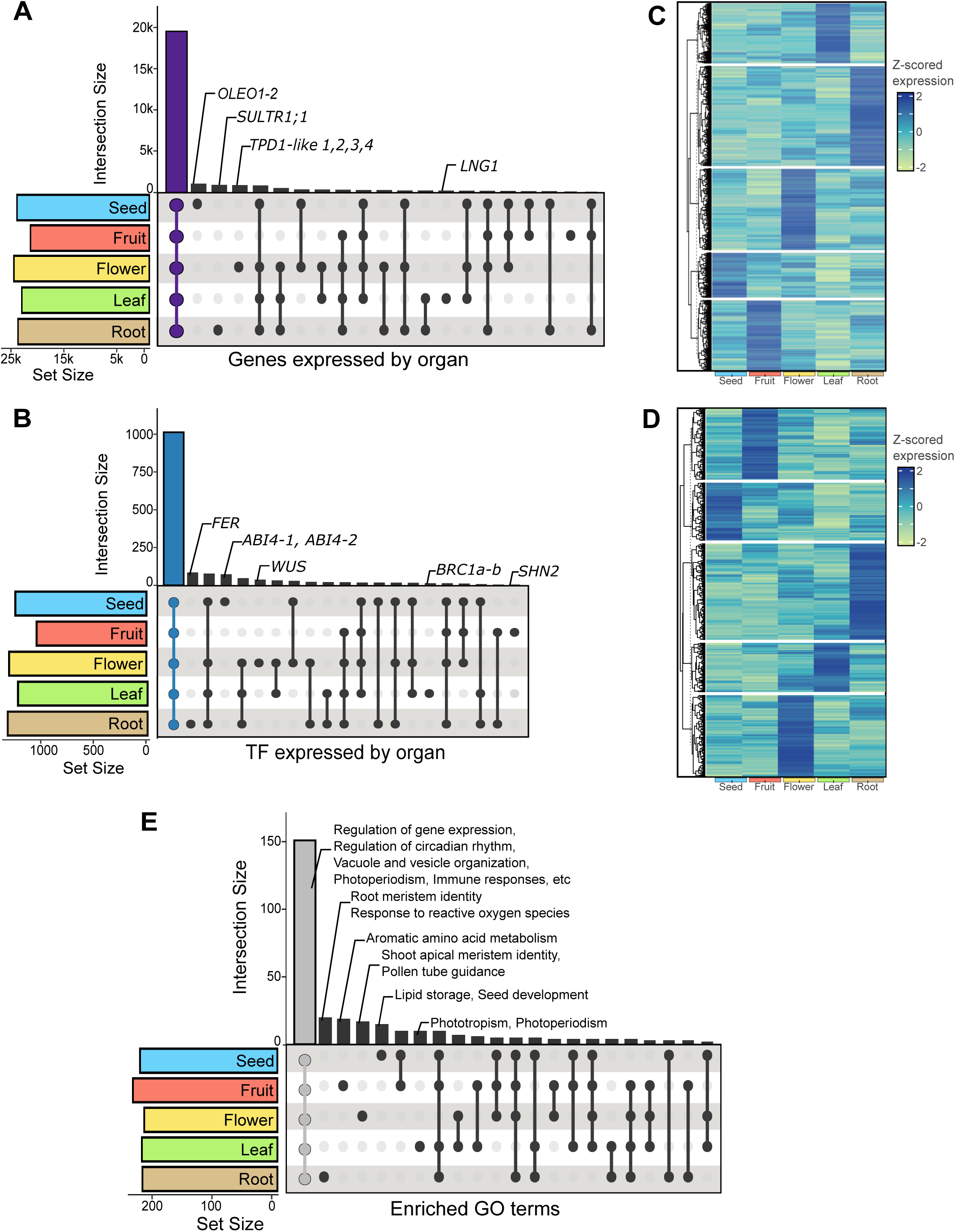
Organ-Level transcriptomic landscape in tomato. Organ-Level transcriptomic landscape in tomato. (A) Distribution of expressed genes across organs. (B) Distribution of expressed transcription factors (TFs) across organs. (C) Heatmap of normalized (Z-scored) expression levels of shared genes across organs. (D) Heatmap of normalized (Z-scored) expression levels of shared TFs across organs. (E) Enriched biological process Gene Ontology (GO) terms associated with expressed genes across organs (FDR-adjusted p-value < 0.05).

**Fig. 2.**
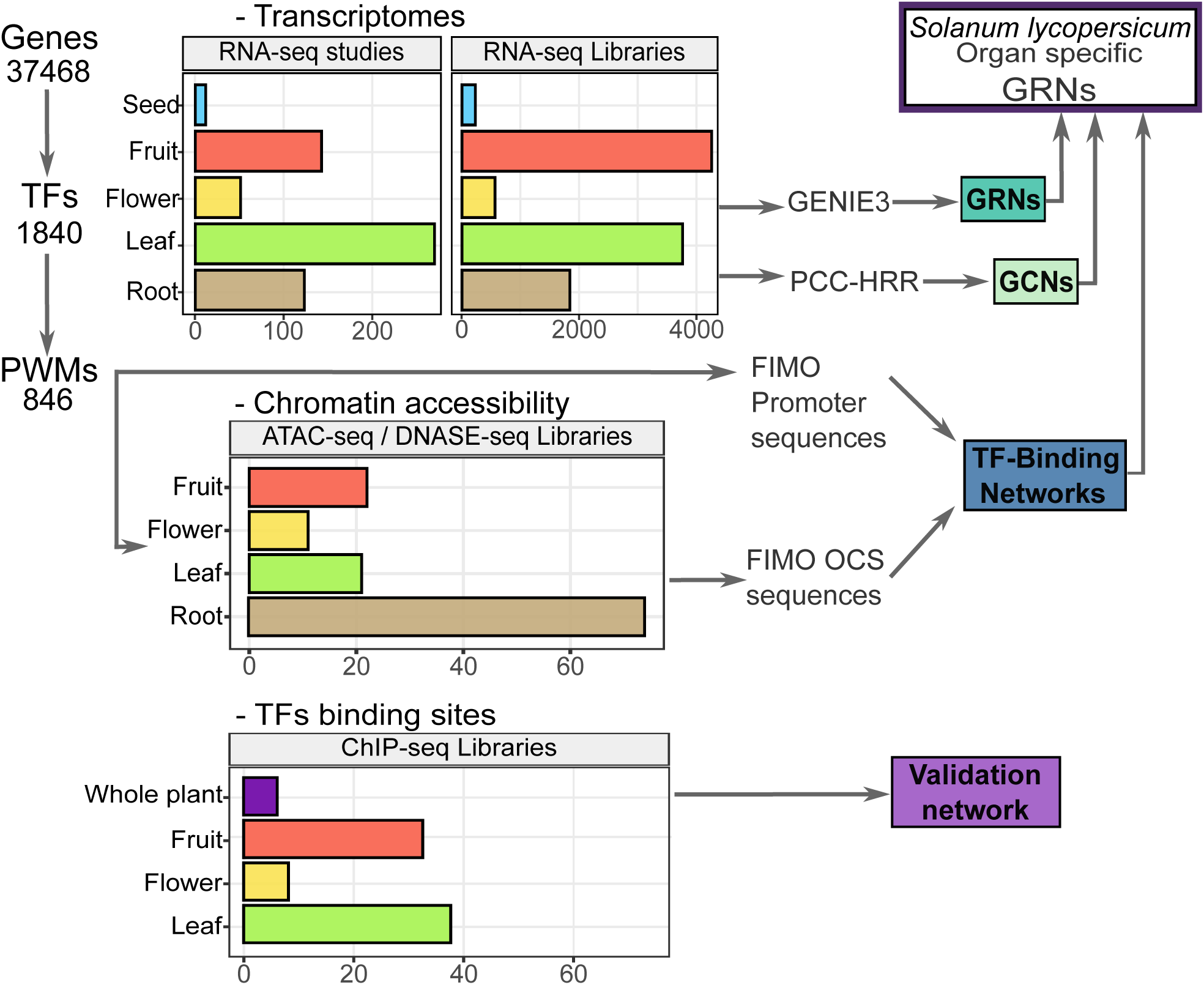
Multi-omics data integration for generating organ-level GRNs in tomato. Overview of datasets used to generate organ-specific gene regulatory networks (GRNs) in *Solanum lycopersicum*. Bar plots indicate the number of available datasets per organ for transcriptomics (RNA-seq), chromatin accessibility (ATAC-seq/DNase-seq), and transcription factor binding sites (ChIP-seq). Arrows illustrate data flow into regulatory network generation, including GRNs, co-expression networks (GCNs), TF-binding networks, and validation datasets.

Concerning TFs, 1612 (87.7%) were expressed in at least one organ. Among them, 1014 (62.9%) were expressed across all organs, while a smaller subset (16.46% of expressed TFs) exhibited organ-level expression (Fig. 1B, Supplementary Table S5). This latter group includes *SlBRC1a* and *SlBRC1b* (*Solyc03g119770* and *Solyc06g069240*), paralogs of Arabidopsis *BRANCHED1*, involved in leaf and axillary bud development (Martín-Trillo *et al*., 2011), *SlFER* (*Solyc06g051550*) a key regulator of root iron uptake (Aviña-Padilla *et al*., 2023), *SlWUSCHEL (WUS, Solyc02g083950)*, which controls floral meristem identity and development (Hawar *et al*., 2022), *SlSHINE2* (*SHN2*, *Solyc12g009490*), encoding a TF that controls epidermal growth in developing fruits (Bres *et al*., 2022), and two Arabidopsis *ABI4* paralogs (*Solyc03g095977* and *Solyc03g095973*) that are exclusively expressed in seeds, consistent with their role in seed vigor (Bizouerne *et al*., 2021). Importantly, while most TFs and other genes are expressed across all organs, their expression levels varied substantially depending on the organ analyzed (Fig. 1C, D). These quantitative differences suggest organ-level regulatory mechanisms, where distinct expression patterns contribute to the specialized functions and characteristics of each organ.

To assess how gene expression in tomato organs relates to the occurrence of organ-specific and widely expressed biological processes, we performed a Gene Set Enrichment Analysis (GSEA) for each organ. The analysis revealed that most enriched biological processes (151 GO terms) were common across all organs (adjusted p-value < 0.05), including gene expression regulation, circadian rhythm, vacuole and vesicle organization, immune responses, mRNA methylation and response to abscisic acid (Figure 1E, Supplementary Table S6). In contrast, only 22 enriched GO terms were identified as unique for each organ. These include processes related to fruit ripening in fruits, root meristem identity and response to reactive oxygen species in roots, phototropism and photoperiodism in leaves, shoot apical meristem identity, brassinosteroid signaling, and pollen tube guidance in flowers as well as lipid storage and seed development in seeds (Figure 1E, Supplementary Fig. S2, Supplementary Table S6). These findings demonstrate that while fundamental biological processes are conserved across all tomato organs, a subset of organ-level processes support their unique functions, emphasizing a complex regulatory mechanism that defines organ identity in tomato.

### Organ-level GENIE3 networks recapitulate experimentally obtained TF-target interactions

To generate organ-level GRNs, we employed the GENIE3 algorithm using the transcriptomic count tables separated by organ and the updated TF list as input. GENIE3 generated a ranked list of putative TF-target interactions, from which we selected the top 1%, 2%, 5%, 8% and 10% of the highest-scoring interactions to evaluate network accuracy. The inferred organ-level networks were benchmarked against ChIP-seq datasets for tomato (Supplementary Table S7). These included data for GLK1 and 2 (Solyc07g053630, Solyc10g008160) (Tu *et al*., 2022), MYC2 (Solyc08g076930) (Du *et al*., 2017), JMJ4 (Solyc08g076390) (Ding *et al*., 2022), WOX13 (Solyc02g082670) (Jiang *et al*., 2023), EIL4 (Solyc06g073730), TAGL1 (Solyc07g055920) and RIN (Solyc05g012020) (Fujisawa *et al*., 2011; Gao *et al*., 2019) (Supplementary Table S7). Each GENIE3 network was benchmarked against a ChIP-seq gold-standard network through Fisher’s exact test enrichment analysis. The analysis included the TFs GLK1 and 2, MYC2, EIL4, JMJ4, and WOX13 to evaluate all organs, while RIN and TAGL1 were assessed exclusively in reproductive organs (excluding roots and leaves networks), as their expression is restricted to these tissues. The top 2% networks showed the highest enrichment (indicated by higher log_2_-transformed Fisher odds ratios) and statistical significance (lower log_10_-transformed adjusted p-values) in the overlap between TF-target pairs obtained by GENIE3 and TF-target pairs obtained by ChIP-Seq (Table 1, Supplementary Table S8). Furthermore, comparison with existing tomato gene networks from PlantRegmap (Tian *et al*., 2020) and TomatoNet (Kim *et al*., 2017) demonstrated that the GENIE3-derived GRNs exhibited greater enrichment and overlap with the gold-standard dataset, indicating a better performance in predicting TF-target interactions obtained experimentally (Supplementary Table S9).

**Table 1.** Enrichment metrics for organ-Level GRNs. Summary of enrichment metrics for organ-specific GRNs, considering the top 2% of interactions identified by the GENIE3 algorithm. Enrichment results obtained from a Fisher’s exact test that assessed gene set overlap significance, reporting log2 fold change, p-value, and intersection size to the validation network (ChIp-seq). -Inf represents log₁₀ adjusted p-values < -400.

|  | Root | Leaf | Flower | Fruit | Seed |
| --- | --- | --- | --- | --- | --- |
| Total TFs | 1,297 | 1,216 | 1,300 | 1,058 | 1,241 |
| Total Genes | 23,226 | 22,513 | 23,988 | 20,888 | 23,124 |
| Total edges | 797,120 | 743,902 | 798,851 | 665,733 | 817,504 |
| log<sub>2</sub> Fisher odd ratio | 1.89 | 2.66 | 2.01 | 3.01 | 2.22 |
| log<sub>10</sub> adj p-value | -156.72 | -inf | -196.33 | -305.17 | -inf |
| Genes in Overlap | 1714 | 2967 | 1912 | 2226 | 4204 |
| AUROC | 0.72 | 0.65 | 0.65 | 0.72 | 0.53 |
| AUPR | 0.51 | 0.45 | 0.46 | 0.52 | 0.32 |

To further evaluate the performance of the top 2% GRNs, we calculated the AUROC (Area under the receiver Operating Characteristic) and AUPR (Area under the precision-recall) curves for each organ-level network and compared these values against a network consisting of TF-target interactions obtained from the gold-standard dataset. The GRNs derived from tomato roots, leaves, flowers, fruits and seeds showed statistically higher AUROC and AUPR values than randomly generated TF-target pairs (Supplementary Fig. S3). These results confirm that the GRNs successfully recapitulate experimentally validated TF-target interactions, underscoring their utility in predicting regulatory interactions for TFs lacking experimental validation.

To provide further support to our networks, we integrated additional layers of information from complementary approaches to the GENIE3-inferred edges. Gene co-expression is widely used to infer biologically relevant relationships between genes (Wolfe *et al*., 2005; Yin *et al*., 2021). Using the same RNA-seq datasets, we generated aggregated gene co-expression networks (GCNs) following the protocol in Orduña et al., (2023). The TF-target pairs from each GCN were extracted as additional evidence for the GENIE3-predicted interactions. While the GENIE3 algorithm predicts regulatory interactions based on expression patterns, additional evidence is necessary to determine whether these interactions occur via direct TF binding to regulatory sequences. To integrate TF-binding information into the GENIE3 networks, we extracted upstream sequences (2 Kb from the transcription start site) for each annotated gene on ITAG4-merge and performed TF-binding motif prediction using the FIMO tool (Grant *et al*., 2011). Additionally, we conducted the same analysis on sequences within open chromatin sites (OCSs) identified in tomato fruit, flower, leaf, and root organs, using data from DNase-seq and ATAC-seq experiments (Supplementary Table S10). The results from all evidence layers were compiled, ensuring that our analysis remained constrained to TF-target pairs identified by GENIE3. This approach maintained the predefined network structure, with additional regulatory evidence mapped onto it rather than introducing new interactions. We found that between 51% and 61% of GENIE3-predicted interactions were supported by at least one additional evidence, with most edges validated by one or two different approaches (Fig. 3A). Furthermore, a substantial portion of GENIE3 edges were validated by the presence of cis-binding motifs detected by FIMO within promoter sequences or OCSs, suggesting a direct regulation of TFs over their inferred targets.

**Fig. 3.**
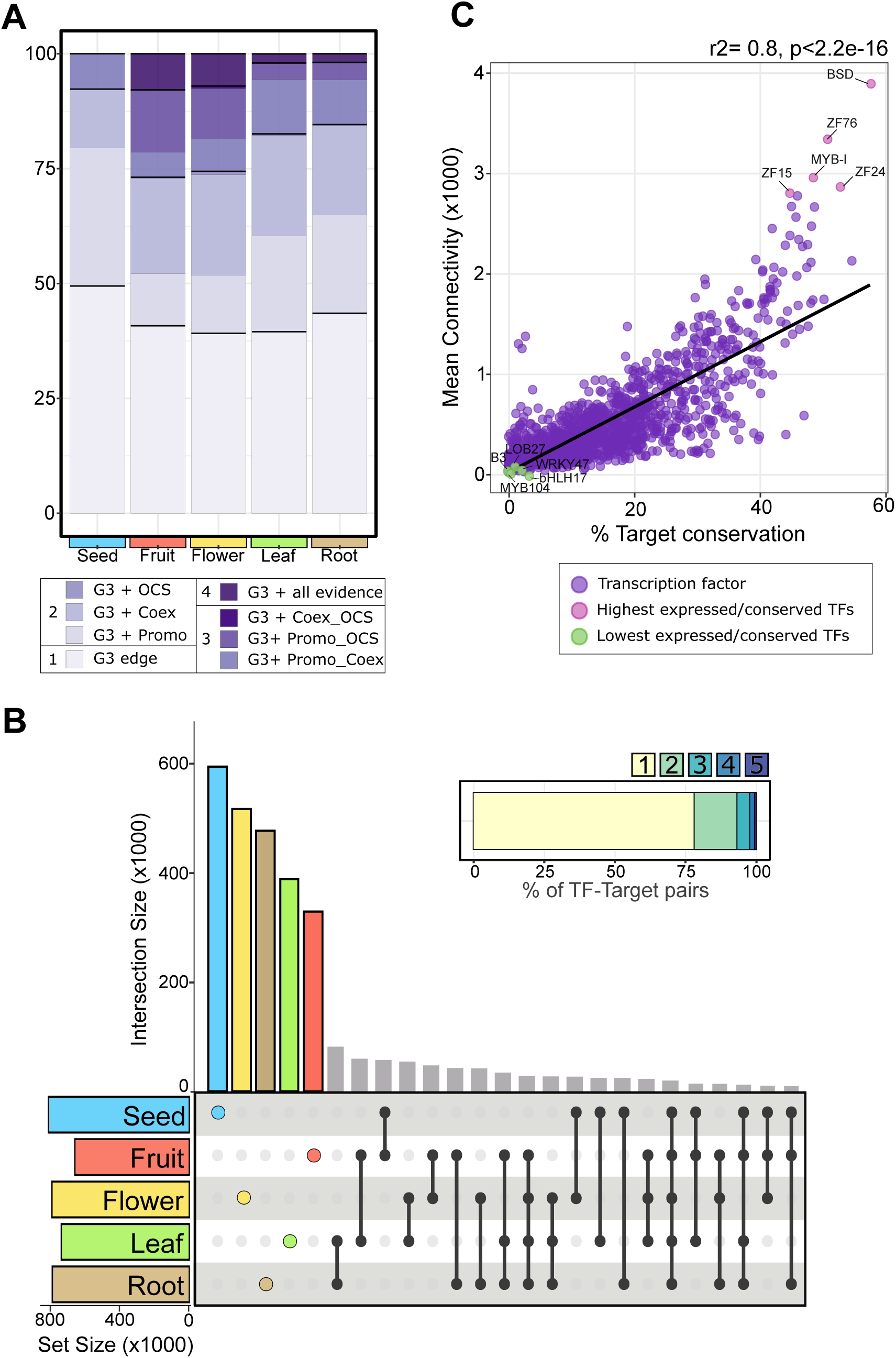
Comparative analysis of organ-level GRNs reveals regulatory signatures and TF connectivity patterns. (A) Stacked bar plot showing the proportion of TF-target pairs in the GENIE3-inferred GRNs supported by multiple evidence sources. Darker purple shades indicate interactions validated by GENIE3 alone (G3) or by 2–4 sources, including open chromatin site binding (OCS), gene co-expression networks (Coex), and promoter binding (Promo). (B) UpSet plot displaying the overlap of TF-target interactions across organ-specific GRNs, with an inset showing the distribution of shared versus unique interactions (1 to 5 organs). (C) Relationship between TF mean connectivity (average number of target genes across organs) and target conservation (proportion of shared targets across organs). Colored dots represent TFs with highest and lowest values. A black trend line highlights the general pattern in the data.

As previously mentioned, most genes and TFs were expressed across all tomato organs, albeit with varying expression levels. To assess how these expression patterns influence regulatory interactions, we examined the distribution of TF-target gene pairs across the five organ-level GRNs. Over 75% of these pairs were unique to a single organ (Fig. 3B), indicating that while TFs and targets are broadly expressed, the existence of regulatory pairs is lar |gely organ specific. This has previously been described for other plant GRNs (Huang *et al*., 2018; Ranjan *et al*., 2024).

To further evaluate how specific TF-target pairs distribute across organs and how target conservation correlates with TF connectivity, we calculated the percentage of conserved targets for each TF across all organs. As expected, most TFs showed low target conservation due to the organ-level nature of TF-target interactions. Interestingly, we observed a significant correlation between TF connectivity and target conservation (R² = 0.36, p < 2.2e^−16^), indicating that highly connected TFs tend to regulate conserved target genes across organs (Fig. 3C; Supplementary Table S11). Gene Set Enrichment Analysis (GSEA) of the top five most connected and conserved TFs revealed strong enrichment for biological process GO terms related to fundamental cellular processes, including nucleic acid metabolism, vesicle transport, and RNA metabolism (FDR-adjusted p-value < 0.05) (Supplementary Fig. S4). These results suggest that TFs with high connectivity and conserved targets function as global regulators of essential cellular pathways. In contrast, TFs with lower connectivity and limited target conservation appear to mediate more specific functions. For instance, despite their ubiquitous expression, TFs such as *LOB27* (*Solyc06g062630*) and *bHLH17* (*Solyc02g093280*) show enriched functions in flowers, *WRKY47* (*Solyc01g058540*) in roots, *B3* (*Solyc05g004000*) in flowers and seeds, and *MYB104* (*Solyc01g090530*) exhibits distinct functions across multiple organs (Supplementary Fig. S5). In sum, organ-level GENIE3 networks present topological features expected for GRNs and can recapitulate TF-target interactions experimentally obtained by complementary approaches.

### The fruit GRN captures known regulatory interactions and identifies novel central controllers of ripening

To evaluate the ability of the organ-level GRNs to capture biologically relevant regulatory interactions, we focused on ripening, one the most extensively studied processes in tomato, linked to hormonal signaling pathways such as ethylene, cell wall remodeling, pigments biosynthesis and other processes (Karlova *et al*., 2014; Li *et al*., 2019; Zhu *et al*., 2022). Tomato fruit ripening is governed by a complex regulatory cascade, involving interactions between well-characterized TFs including APETALA2a (AP2a), NON-RIPENING (NOR), FRUITFULL (FUL1/TDR4 and FUL2/MBP7), TOMATO AGAMOUS-LIKE 1 (TAGL1), RIPENING INHIBITOR (RIN) and COLORLESS NON-RIPENING (CNR) (Li *et al*., 2021a; Zhu *et al*., 2022). Among these, TAGL1 and RIN are recognized as central regulators of ripening (Gao *et al*., 2019; Li *et al*., 2021b).

To determine whether the fruit GRN reproduced regulatory interactions of known TFs, we compared the targets of TAGL1 and RIN from the fruit GRN with gene lists compiled from previous omics studies. These included differentially expressed genes (DEGs) identified in TAGL1 and RIN knockout and RNAi plants (Gao et al., 2019; Ito et al., 2020; Li et al., 2018), as well as direct binding targets identified via RIN ChIP-chip and ChIP-seq (Fujisawa et al., 2013; Gao et al., 2019; Zhong et al., 2013), and TAGL1 ChIP-seq (Gao *et al*., 2019) experiments. For RIN, we observed a statistically significant overlap of the targets determined in the fruit GRN with targets obtained in all the experiments, including ChIP-binding targets and regulatory targets identified in RIN-deficient plants (Fig. 4A). Similarly, for TAGL1, the fruit GRN was able to identify targets validated by ChIP-seq and/or knock-out experiments (Fig. 4B). This finding highlights the potential of the GENIE3 GRN to capture experimentally validated regulatory interactions, with many of these corresponding to direct binding of a TF to a target promoter (45% of RIN GRN targets (411/857) and 89% of TAGL1 GRN targets (740/827) are validated by ChIP binding evidence) (Figure 4A-B).

**Fig. 4.**
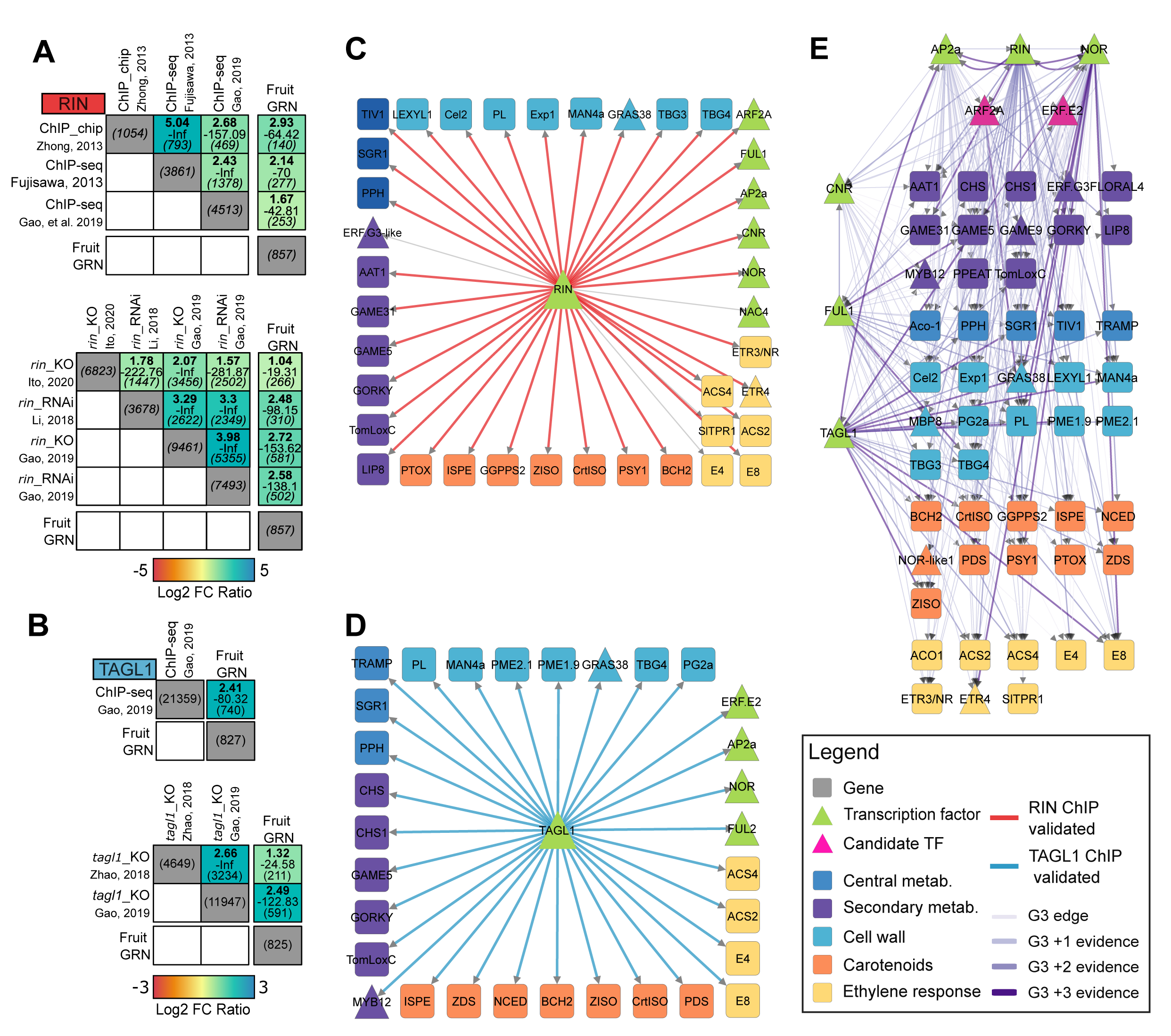
Identification of key transcriptional regulators of fruit ripening regulatory cascades in tomato. (A–B) Enrichment and validation of fruit GRN for RIN (A) and TAGL1 (B) using knockout mutant data and ChIP-binding analyses. Box heatmaps display enrichment obtained from a Fisher’s exact test (log2 fold change, p-value, and intersection size) to the validation network (ChIp-binding). (C–D) Network representation of ripening-associated genes (Li et al., 2019; Zhu et al., 2022) for RIN (C) and TAGL1 (D) derived from the fruit GRN. Triangles represent TFs, squares represent target genes. Node colors indicate function. Edges are colored in red (RIN) or blue (TAGL1) when interactions are validated by ChIP-binding evidence. (E) Network of key regulators of ripening-associated genes (Top scored TFs), applying the same node and edge color scheme as in (C–D). Edge darker shades indicate accumulated regulatory evidence.

To further investigate the regulatory roles of RIN and TAGL1 in fruit ripening, we generated RIN and TAGL1 subnetworks focusing on genes involved in fruit ripening described in Li *et al*., (2019) and Zhu *et al*., (2022). We found that the GENIE3 network predicts interactions for both RIN and TAGL1 with an important number of ripening genes (Supplementary Table S12). Furthermore, most of the GENIE3 edges connecting RIN and TAGL1 to these target genes (90% for RIN and 100% for TAGL1) are supported by ChIP evidence (Fig. 4C, D). Additionally, the subnetworks highlight the regulatory influence of RIN and TAGL1 across diverse biological processes including central or secondary metabolism, cell wall modification, and their interactions with other known TFs in fruit ripening including CNR, NOR, and AP2a (Fig. 4C, D).

To identify novel ripening regulators, we used the fruit GENIE3 network to generate a subnetwork including the list of ripening-associated genes (Li *et al*., 2019; Zhu *et al*., 2022). We applied the Integrated Value of Influence (IVI), a metric that combines centrality measures such as degree centrality, cluster rank, neighborhood connectivity, betweenness centrality, and collective influence into a single value to quantify network hub influence (Salavaty *et al*., 2020). We found that among TFs with higher IVI are CNR, NOR, FUL1, AP2a, RIN, and TAGL1, confirming their roles as central regulators of fruit ripening genes. Notably, two additional TFs, *Sl*ARF2A (Solyc03g118290) and *Sl*ERF.E2 (Solyc06g063070), emerged as TFs with high IVI, revealing a potentially relevant role in controlling ripening-related genes (Supplementary Table S13). *Sl*ARF2A TF has been identified as a regulator of axillary shoot development and is predominantly expressed in the late stages of ripening (Xu *et al*., 2016). RNAi lines targeting *Sl*ARF2A exhibit ripening defects and ethylene insensitivity, while overexpression lines show accelerated and uneven ripening (Hao *et al*., 2015; Breitel *et al*., 2016). On the other hand, *Sl*ERF.E2 expression is downregulated in *cnr*, *nor*, and *rin* mutants, however its role in ripening has not been elucidated yet (M. Liu et al., 2016). Interestingly, we found that *Sl*ARF2A and *Sl*ERF.E2 may act upstream of important TFs such as AP2, NOR, and CNR, in addition to multiple ripening-relevant genes (Fig. 4E).

### Tomato organ-level GRNs validate the role of ABF TFs on abscisic acid (ABA) regulatory cascades and identify SlGBF3 as a new regulator of ABA-related genes

ABA is a key plant hormone involved in the regulation of seed dormancy, germination, seedling development, root growth, flowering, and responses to abiotic and biotic stresses (Vishwakarma *et al*., 2017; Krukowski *et al*., 2023). Notably, the GO term “response to ABA” was consistently enriched across all organs (Supplementary Fig. S2). This category includes 730 genes, 714 of which are ubiquitously expressed in tomato organs (Supplementary Table S14, hereafter referred as “ABA-related genes”). Among known TFs participating in ABA responses are the members of the AREB/ABF (ABA response element binding/ABA response element binding factor) family of bZIP TFs (Uno *et al*., 2000; Krukowski *et al*., 2023). This family has ten members in tomato (Pan *et al*., 2023), however the relative contribution of each ABF TF at the organ-level remains unexplored. To address this, we constructed organ-level networks for the ten ABFs, focusing specifically on their regulation of ABA-related genes. While most ABF family members are expressed at similar levels across organs, -except for *Sl*ABF6 (not expressed in flowers) and *Sl*ABF7 (not expressed in fruits and leaves), their regulatory potential differs depending on the organ. *Sl*ABF1 and *Sl*ABF4 appear to play key roles in regulating ABA-related genes in fruits, whereas *Sl*ABF2, *Sl*ABF3, *Sl*ABF5, and *Sl*ABF10 regulate more genes in the leaf network. *Sl*ABF5, *Sl*ABF9, and SlABF10 are involved in ABA response regulation in roots, while *Sl*ABF2 seems to play a more important role in the flower GRN. For seeds, *Sl*ABF6 and *Sl*ABF7 exhibit the highest regulatory activity on ABA-related genes (Figure 5A, Supplementary Table S15). These findings point out that the ABF TFs have different relative contributions on the regulation of ABA-related genes across organs.

**Fig. 5.**
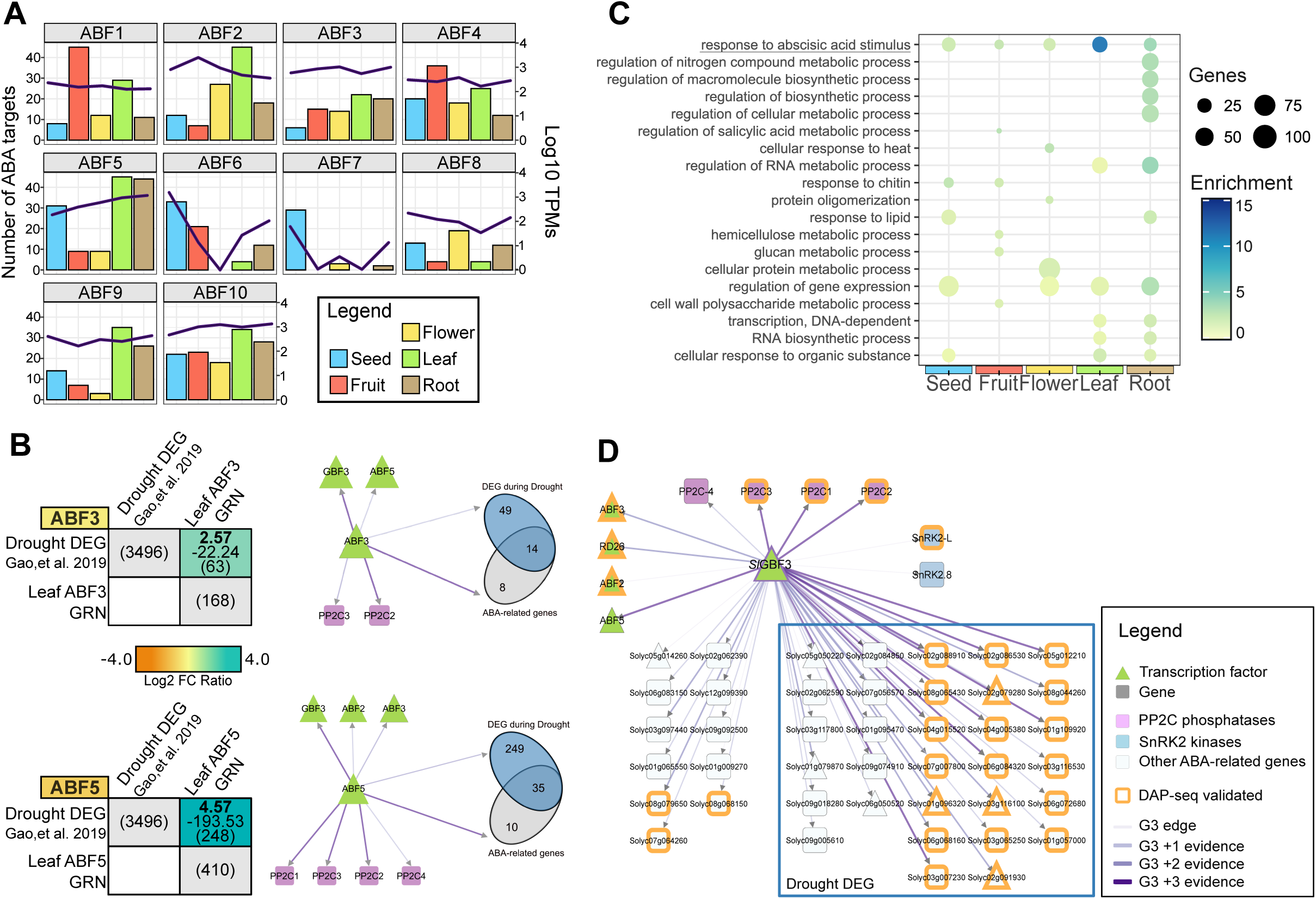
Tomato GRNs reveal the role of ABF TFs and identify a key regulator of ABA-related GRNs. (A) Bar plots showing organ-specific expression levels and the number of ABA-related target genes regulated by ABF TFs. Bars indicate target counts, while black lines represent log₁₀ TPM expression values. (B) Enrichment and validation of leaf GRNs for *ABF3* and *ABF5* using drought-responsive DEGs (Gao et al., 2019). Box heatmaps (left) display enrichment results from a Fisher’s exact test (log2 fold change, p-value, and intersection size). The networks (right) show distribution of ABA-related and drought regulated targets of these TFs. (C) Gene Set Enrichment Analysis (GSEA, FDR-adjusted p-value < 0.05) of *SlGBF3* target genes in organ-specific GRNs. Dot size represents gene number, while color intensity reflects enrichment values. (D) Network visualization of *SlGBF3*-regulated ABA-related genes in the leaf GRN. Triangles represent TFs, squares represent target genes. Node colors indicate function. Orange-bordered nodes indicate DAP-seq validated targets, while enclosed nodes are DEGs from drought-stressed leaves (Gao et al., 2019). Edge darker shades indicate accumulated regulatory evidence, (GENIE3: G3).

Plant response to drought is closely linked to ABA signaling (Kang *et al*., 2002; Krukowski *et al*., 2023). To validate the regulatory predictions from our GRNs, we focused on two TFs, *Sl*ABF3 and *Sl*ABF5, which have been identified as relevant regulators of drought responses in tomato leaves (Kang *et al*., 2002; Hsieh *et al*., 2010). Using the leaf GRN, we extracted the predicted targets of ABA-related genes from both TFs and compared them to the DEGs found in a transcriptome analysis of leaves from drought-exposed plants (Wang et al., 2023). Our analysis revealed a significant enrichment of drought-responsive genes among the targets of both TFs (Fig. 5B, Supplementary Table S16). We found that 284 out of 410 (∼69%) ABF5 targets and 63 out of 168 (∼38%) ABF3 targets overlapped with drought-responsive DEGs. Additionally, 14 out of 22 (∼64%) ABF3 targets and 35 out of 45 (∼78%) ABA-relevant genes were also present among the drought DEGs. The GRN also indicated that both TFs regulate ABA signaling components encoded by *protein phosphatase class 2 C (PP2C)* genes (Kang *et al*., 2002; Fujii *et al*., 2009; Krukowski *et al*., 2023), including *Solyc03g121880* and *Solyc05g052980* (regulated by both TFs), and *Solyc03g096670* and *Solyc06g076400* (regulated specifically by *Sl*ABF5). Furthermore, we observed a potential feedback regulatory mechanism between *Sl*ABF3 and *Sl*ABF5, as previously reported for Arabidopsis (Chang *et al*., 2019) (Fig. 5B). These findings confirm the key roles of *Sl*ABF3 and *Sl*ABF5 in the ABA-mediated drought regulatory cascades (Kang *et al*., 2002; Hsieh *et al*., 2010; Krukowski *et al*., 2023; Pan *et al*., 2023).

To identify novel regulators of ABA responses beyond the ABF family of TFs across all organs, we filtered the five organ-level GRNs to retain TFs with regulatory connections to ABA-related genes. A network analysis calculating the Integrated Value of Influence (IVI) (Salavaty *et al*., 2020) of the network hubs identified the *Sl*GBF3 TF (Solyc01g095460) as one of the top ten most influential TFs in the ABA-related networks across all organs, and the most influential in the leaf GRN (Supplementary Table S17). The *Sl*GBF3 has recently been found to be co-expressed with drought-responsive genes in tomato leaves (Bortolami *et al*., 2024). To further explore the role of *Sl*GBF3, we performed a GSEA on its target genes in each organ-level GRN. We found a significant enrichment of genes belonging to the “response to abscisic acid stimulus” biological process shared across all organ networks, indicating a potential conserved role in the regulation of ABA-related genes (Fig. 5C).

To further validate the potential regulatory interactions of *Sl*GBF3 identified in the GRNs, we performed DNA Affinity Purification Sequencing (DAP-seq). The DAP-seq analysis identified approximately 46,705 binding events (peaks) associated with genomic regions. These peaks were linked to a total of 11,381 genes annotated in ITAG4-merge in both experimental replicates (Supplementary Table S18, Supplementary Fig. S6A). Using the identified peak regions, we predicted the DNA-binding motif of *SlGBF3* through a MEME-chip analysis (Machanick and Bailey, 2011). Our analysis revealed that *Sl*GBF3 predominantly binds to a consensus sequence “AYGTGGCA” (e-value = 5.5 e^-132^ from 28,413 sites) (Fig. S6B). Comparative analysis with existing TF binding motifs in Cis-BP (Weirauch *et al*., 2014) and JASPAR (Castro-Mondragon *et al*., 2022) databases for *Arabidopsis thaliana* GBF3 (*At*GBF3) showed that it has three binding motifs (MA1351.1-3) with an average consensus sequence of “CGTGGCA” (Fig. S6C). Notably, the binding motif of *Sl*GBF3 exhibited a significant correlation (Pearson correlation coefficient between 0.7-0.865) against *At*GBF3 motifs (Fig. S6D), indicating a strong evolutionary conservation between these TFs and validating our experimental results.

To evaluate the functional relevance of the identified binding events, we compared the predicted targets in the tomato organ-specific GRNs with those identified through DAP-seq. Genes that overlapped between these datasets were classified as high-confidence targets (HCTs). Our analysis revealed that 42–58% of the GRN-predicted targets of *SlGBF3* were supported by DAP-seq binding evidence, with this overlap being significantly enriched (Supplementary Fig. S7). A GSEA of the complete network of HCTs according to DAP-seq revealed a significant enrichment in functions related to water deprivation, abiotic stimuli, and hormone responses, with ABA as a central component (Supplementary Fig. S8). To further explore *Sl*GBF3 role in ABA-related regulatory networks, we constructed GRNs specifically for ABA-responsive genes across different organs. Among these, the leaf GRN exhibited the highest representation of ABA-related genes. Notably, over 60% of the targets of *Sl*GBF3 in the leaf network were associated with drought stress responses, including multiple *PP2C* genes (*Solyc03g121880*, *Solyc03g096670*, *Solyc05g052980*, *Solyc06g076400*) and two *SNF1-related protein kinase 2 (SnRK2)* genes (*Solyc08g077780*, *Solyc04g012160*) (Fig. 5D). Additionally, *Sl*GBF3 appears to function as an upstream regulator of key TFs controlling ABA-related genes, including *SlABF2*, *SlABF3*, and *SlABF5*. In the leaf network, over 50% of ABA-related targets were classified as high-confidence targets (HCTs) based on DAP-seq evidence (Fig. 5D). In ABA-related GRNs from other organs, *Sl*GBF3 was found to regulate smaller subsets of ABA-responsive genes, including *PP2C* genes, ABF TFs, and key regulators such as MYB1 (Solyc12g099120), a TF implicated in ABA-mediated pathogen susceptibility (Abuqamar *et al*., 2009) (Supplementary Fig. S10). Although the leaf-specific GRN contained the largest number of ABA-related targets, all organ-specific ABA networks maintained a consistent proportion of HCTs (∼50%) (Supplementary Fig. S9). This result suggests a conserved regulatory role of *Sl*GBF3 across different organs, reinforcing its significance in ABA-mediated stress responses.

### TomViz-GRN App: An online tool for the visualization of tomato GRNs

To provide the scientific community with a comprehensive framework of tomato organ-level GRNs and a user-friendly resource, we have developed a public web platform featuring an interactive interface that allows users to explore the results of this study. Our GRN apps within the TomViz module of the PlantaeViz platform (https://plantaeviz.tomsbiolab.com/tomviz) (Santiago *et al*., 2024) adhere to the Findability, Accessibility, Interoperability, and Reusability (FAIR) principles. Through the website app, users can interact with organ-level GRNs, select and subset network data for download (Fig. 6A, C). The TomViz-GRNs app provides various features for data analysis. In the Regulatory Targets Tab, users can query individual TFs or genes to explore central regulatory TFs and their validation layers (Fig. 6B). The D3 Subnetwork Tab allows users to upload gene lists and generate GRNs based on specific queries. It categorizes data and enables an organ-level study of stress responses, helping to detect novel regulatory pathways and TFs involved in multiple regulatory cascades (Fig. 6C). The TomViz-GRNs app thus provides an intuitive platform for studying tomato gene regulation and investigating stress responses across different organs.

**Fig. 6.**
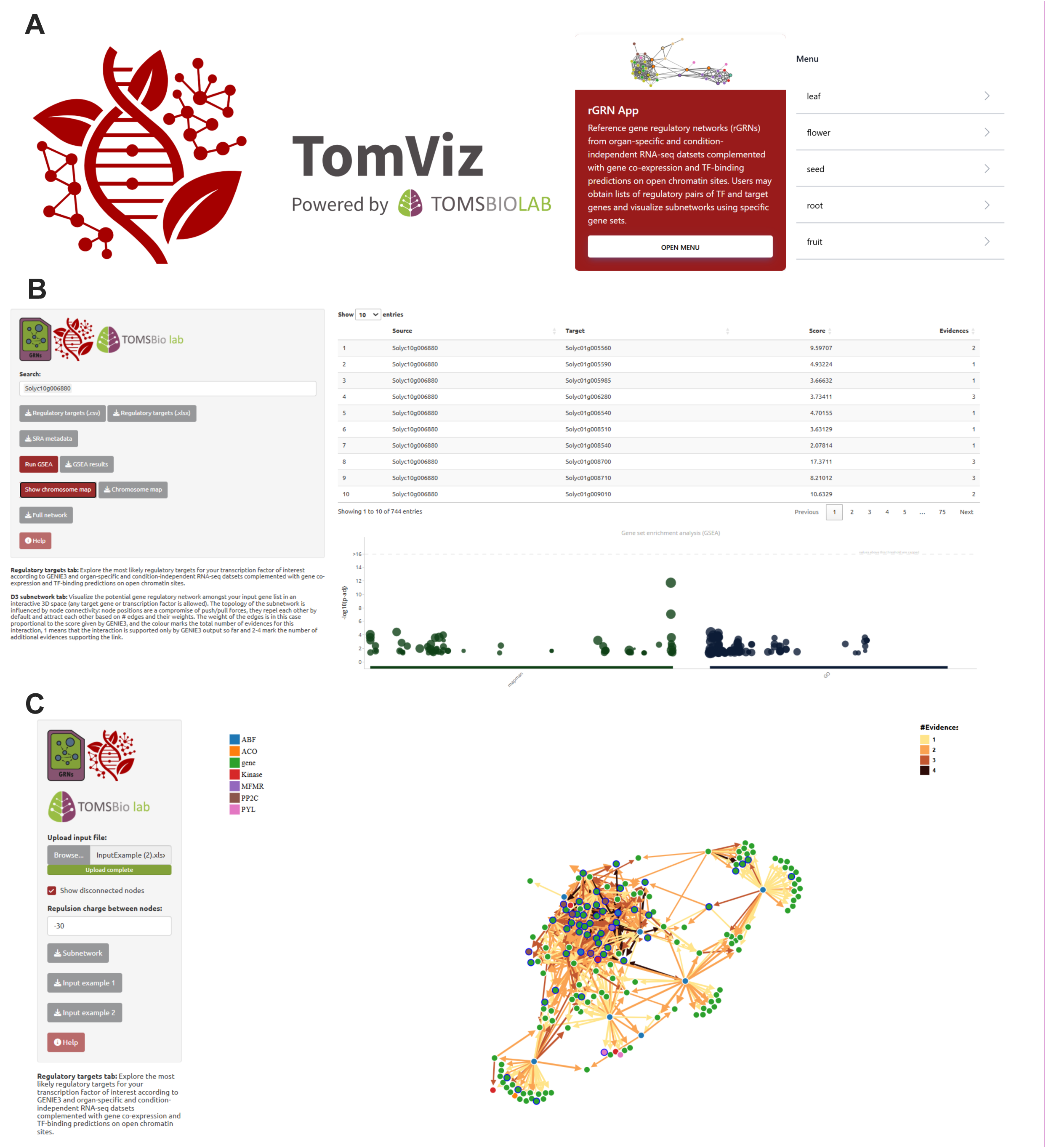
TomViz-GRNs: A web-based platform for exploring tomato organ-level GRNs. (A) TomViz interface within the PlantaeViz platform (Santiago et al., 2024), providing access to GRN exploration tools. (B) Regulatory Targets Tab: Users can query TFs or genes to explore regulatory interactions and validation layers. The interface includes options to download TF target lists, perform GSEA, and visualize target distributions on a chromosome map. (C) D3 Subnetwork Tab: Users can upload gene lists, visualize GRNs, and analyze regulatory pathways at the organ level. The visualization includes directional edges representing regulatory interactions from TFs to targets, with edge colors indicating the level of supporting evidence. Additional options allow customization of network layout and node separation.

## Discussion

### Bioinformatic validation of GRNs

To generate tomato GRNs, our first step was to generate an updated resource of gene models, TFs and functional annotations. Using well-established gene functional annotation pipelines (Jones *et al*., 2014; Cantalapiedra *et al*., 2021), we successfully assigned functional annotations to 26,403 of the 37,467 ITAG4-merge genes, considerably improving coverage compared to the previous ITAG4.1 annotation in SolGenomics, which included only 13,142 functionally annotated genes or previous annotations for iTAG4.0 (25,285 genes) (Rivera-Silva *et al*., 2024).

TF prediction remains a challenging task, as automated approaches usually rely on protein sequence scanning for known DNA-binding domains, which may lead to the inclusion of proteins with DNA-binding capabilities unrelated to TF function (Itzkovitz *et al*., 2006; Liebold *et al*., 2024). To avoid such proteins and refine the tomato TF list, we integrated multiple levels of evidence to filter and extract a curated set of 1,840 TFs (representing around 5% of tomato genes). This number closely aligns with TF numbers reported for tomato in PlantTFDB (1,845 TFs) (Jin *et al*., 2017) and is slightly higher than those in CisBP (1,773 TFs) (Weirauch *et al*., 2014) and other annotated TFs from previous studies (1,069 TFs in Kumar et al., 2021). Overall, the percentage of TF coding genes we found is slightly lower than TFs reported for Arabidopsis (approximately 5-10%) (Riechmann and Ratcliffe, 2000) but consistent with estimates reported for other crops, including wheat (5.7%) and rice (6.1%) (Zheng *et al*., 2016). It is also comparable to values reported for other Solanaceae species, such as eggplant (5.3%) (Wei *et al*., 2020).

A vast number of transcriptomic studies have been conducted in tomato, covering diverse experimental conditions, organs, and developmental stages. While several efforts have aimed to generate gene expression atlases for tomato gene expression (Ozaki *et al*., 2010; Fukushima *et al*., 2012; Gao *et al*., 2013; Koenig *et al*., 2013; Arhondakis *et al*., 2016; Zouine *et al*., 2017; Bae *et al*., 2021; Bizouerne *et al*., 2021; Kumar *et al*., 2021; Kusano *et al*., 2022; Li *et al*., 2024) many studies are limited in scope, often focusing on specific experimental conditions, using older genome assemblies (SL2.4 or SL3.0 with iTAG2.5 or iTAG3.0 annotations), or relying on microarray data. To our knowledge, the gene expression dataset collected in this study represents the most comprehensive to date, compiling over 10,000 RNA-Seq libraries from five major organs and integrating hundreds of Bioprojects performed worldwide. Moreover, the transcriptomes were processed utilizing the latest genome version (SL4.0) and the updated ITAG4-merge annotation, resulting in greater gene coverage. This extensive dataset enabled us to characterize general gene expression patterns at the organ level, facilitating the identification of genes involved in organ-specific functions.

Consistent with previous studies in tomato (Li *et al*., 2024), we found that most genes meet the threshold for expression across all organs. Similar ubiquitous expression patterns have been reported in other plants, including *Linum usitatissimum* (Qi *et al*., 2023) and *Zea mays* (Huang *et al*., 2018), where over 50% of genes are expressed across multiple tissues. Despite this ubiquitous expression, we found differences in expression levels across organs for these genes, showing that organ identity is an important determinant of differential gene expression, as has been shown in previous studies in Arabidopsis (Aceituno *et al*., 2008).

Beyond broadly expressed genes, we identified a substantial subset of genes with organ-specific expression that were enriched in biological processes critical for organ function. Our analyses successfully captured known organ-specific genes. Moreover, genes such as *SULTR1;1* and *FER* in roots, *TPD1-l* genes in flowers, *LNG1* and *SlBRC1a* in leaves, stems, and hypocotyls, and ABI TF in seeds were also identified as organ-specific in a recent study in tomato (Li *et al*., 2024). Furthermore, the TFs also displayed widespread expression across tomato organs, mirroring findings in *Arabidopsis* (Ranjan *et al*., 2024). Prior studies in tomato indicate that fewer than 20% of expressed TFs are organ-specific (Rohrmann *et al*., 2012). Nonetheless, despite their broad expression, variations in TF expression levels across organs highlight the dynamic and context-dependent regulation of transcriptional networks governing organ function.

Transcriptomic data has been extensively used to generate biological network models with the purpose of identifying key candidates for functional genomics analyses. Due to the limited availability of TF-target interaction data existing for tomato, the majority of studies have relied on GCNs to infer regulatory relationships and identify co-regulated gene groups (Fukushima *et al*., 2012; Koenig *et al*., 2013; Ichihashi *et al*., 2014; Arhondakis *et al*., 2016; Yue *et al*., 2016; Kim *et al*., 2017; Zouine *et al*., 2017; Bizouerne *et al*., 2021; Kusano *et al*., 2022; Pirona *et al*., 2023; Wang *et al*., 2023a). Notably, the GCNs lack directionality, making it difficult to establish regulatory interactions. Additionally, many rely on correlations such as Pearson coefficients, which fail to capture non-linear relationships (Escorcia-Rodríguez *et al*., 2023). To address these limitations, we employed GENIE3, a widely used algorithm for reconstructing directed GRNs in plants *(Chen et al., 2023; De Clercq et al., 2021; Harrington et al., 2020; Huang et al., 2018; Ranjan et al., 2024; Tu et al., 2020)* and other organisms (Huynh-Thu *et al*., 2010; Huynh-Thu and Geurts, 2019; Cuesta-Astroz *et al*., 2021; Olivares-Yañez *et al*., 2021). GENIE3 requires only gene expression data as input, making it particularly suitable for tomato, where TF gene targets remain poorly characterized.

To validate our GRNs, we benchmarked them against available ChIP-Seq data standard networks, employing the AUPR and AUROC curves. This strategy, previously used to assess GRN performance in plants (Brooks *et al*., 2019; Contreras-López *et al*., 2022), provides a more centered evaluation of TF-target interactions than alternative methods that utilize gene co-association to biological processes or metabolic pathways (Kim *et al*., 2017; Orduña *et al*., 2023). Our analysis determined that a 2% best GENIE3 scores threshold—comprising 660,000–800,000 edges—was a suitable cut-off, aligning with previous GENIE3-based GRN studies in crops, which typically consider networks containing around 1 million edges (Harrington et al., 2020; Huang et al., 2018; Ramírez-González et al., 2018). Notably, the GENIE3-derived GRNs outperformed other tomato networks from public resources such as PlantRegMap (Tian *et al*., 2020) and other genome-scale biological network models (Kim *et al*., 2017). Additionally, more than 50% of GENIE3-predicted edges were supported by one or more independent evidence, including cis-regulatory motif binding predictions, further reinforcing the biological relevance of these regulatory connections, as it was proven in other networks (De Clercq *et al*., 2021; Chen *et al*., 2023). This integration of multiple validation strategies enhances the accuracy and functional significance of inferred GRNs, providing a robust framework for studying transcriptional regulation in tomato.

Our analysis revealed that while most genes are broadly expressed across organs, the vast majority of TF-target interactions remain organ-specific. This pattern has been observed in GRNs from other crops (Huang *et al*., 2018; Ranjan *et al*., 2024), suggesting that gene expression levels play a relevant role in establishing regulatory interactions underlying organ-specific functions. Notably, we identified a positive correlation between TF connectivity and target conservation across organs. Highly connected TFs (hubs) tend to regulate a similar set of targets in all organs, whereas TFs with fewer connections are more likely to control organ-specific processes. For example, a highly connected TF *SlHUA* (*Solyc12g017410*) primarily regulates genes involved in fundamental cellular functions, while *SlNGA3* (*Solyc05g004000*) a TF with limited connectivity exhibits distinct, tissue-specific roles—regulating auxin response in flowers and defense responses to fungi in seeds, with no enriched function detected in other organs. Accordingly, *SlNGA3* is homologous to *AtNGA3*, which is associated with flower development (Salava *et al*., 2022). An evolutionary constraint may underlie this phenomenon, as TF-target interactions involving hub genes are more conserved. Disruptions in these interactions are more likely to be deleterious, leading to reduced genetic diversity and slower evolutionary rates among hub TFs. In contrast, tissue-specific TFs, which are less connected, have been described to evolve faster (Mack *et al*., 2019).

### Biological validation of GRNs involved in ripening

We employed the fruit-specific GRN to identify important TF regulators of fruit ripening, including RIN and TAGL1 (Karlova *et al*., 2014). To assess the accuracy of our networks, we compared their predicted TF-target interactions against experimentally validated target lists, which included genes confirmed through TF binding studies (Fujisawa *et al*., 2013; Zhong *et al*., 2013; Gao *et al*., 2019) and regulatory studies of TF-deficient plants (Li *et al*., 2018; Zhao *et al*., 2018; Gao *et al*., 2019; Ito *et al*., 2020). Our results showed a significant enrichment between the predicted interactions in our GRNs and these validated targets, supporting the networks’ ability to accurately capture *in vivo* regulatory relationships. Notably, our networks identified RIN targets such as *FUL1*, previously validated via yeast one-hybrid assays (Fujisawa *et al*., 2014), as well as essential ripening genes, including *ACS2, ACS4, E8, EXP1, PSY, NOR,* PSY and *CNR*, which were confirmed using ChIP-PCR (Martel *et al*., 2011). Similarly, qPCR validation supported TAGL1 targets, including *FUL2* (Fujisawa *et al*., 2014), and ripening-associated genes like *ACS2, ETR1, ERF2* and *PL* (Itkin et al., 2009). These findings underscore the predictive strength of our GRNs in identifying key regulatory interactions governing fruit ripening.

Interestingly, *RIN* and *TAGL1* regulate overlapping sets of ripening-related genes, but our fruit-specific GRN does not predict a direct regulatory link between these two TFs. This suggests that their influence on fruit ripening may be mediated through indirect interactions, such as protein-protein interactions or epistatic synergistic control of ripening-responsive genes, as proposed in previous studies (Fujisawa et al., 2014; Jeon et al., 2024). Notably, our GRN identified *Sl*ARF2A and *Sl*ERF2.E2 as central hubs in the fruit ripening regulatory network, suggesting that these TFs may play crucial roles in modulating the ripening process. *Sl*ARF2A has been implicated in the hormonal regulation of tomato fruit ripening. RNAi-mediated silencing of *Sl*ARF2A results in ripening defects and ethylene insensitivity, while its overexpression (OX) leads to accelerated and uneven ripening (Hao *et al*., 2015; Breitel *et al*., 2016). Consistently, our network analysis identified *Sl*ARF2A targets that align with gene expression changes in OX-ARF2A plants, including *ETR*, *ACS4*, *AP2A*, *ETR3*, *ETR4*, *NOR*, and *RIN* (Breitel *et al*., 2016). In contrast, *Sl*ERF.E2 is associated with key ripening regulators and ripening-related genes, as its expression is downregulated in *cnr*, *nor*, and *rin* mutants, yet its precise function remains unknown (Liu *et al*., 2016). Given their potential regulatory roles, future studies should employ TF-binding assays to validate their direct interactions with ripening-responsive genes and further elucidate their contributions as key hubs of tomato fruit ripening.

To further explore organ-specific regulatory mechanisms, we examined the cellular response to abscisic acid (ABA), a biological process that was ubiquitously enriched in our organ specific gene lists. Our GRN analysis provided new insights into ABA response cascades, particularly in relation to drought-response regulation. We identified *Sl*ABF3 and *Sl*ABF5 as important controllers of drought-responsive genes, as well as ABA-related genes such *as PYL/RCAR transporters*, *PP2Cs phosphatases* and *SnRK2s protein kinases* (Fujii *et al*., 2009). We found *Sl*ABF3 targets *Sl*ABF5 and two *PP2C phosphatases*, a direct interaction previously confirmed in yeast two-hybrid assays (Chen *et al*., 2016). Furthermore, *Sl*ABF5 showed a strong connection to drought responses, as demonstrated by the significant enrichment of its target genes in the drought responsive transcriptome (Wang et al., 2023). Additionally, its expression levels vary during drought stress, as reported in earlier studies (Orellana *et al*., 2010; Wang *et al*., 2023b). These findings confirm that our leaf-specific GRN successfully captures ABA-dependent regulatory pathways, reinforcing the role of *Sl*ABF3/*Sl*ABF5 as regulators of drought-responsive genes expression in tomato leaves.

### SlGBF3 as a novel controller of ABA-related and drought-responsive genes

Our study findings identify the *Sl*GBF3 TF as a novel regulatory hub in ABA-responsive GRNs, suggesting a key relevance in the drought response and ABA signaling cascades across different organs, especially in leaves. Prior research (Bortolami *et al*., 2024) linked *Sl*GBF3 to drought-responsive co-expression modules in tomato, but its direct regulatory function remained unknown. Using DAP-seq analysis, we show direct molecular evidence that *Sl*GBF3 binds to and regulates multiple ABA-related genes, establishing it as a key component of ABA-driven transcriptional regulation in tomato. Importantly, our ortholog and binding motif analyses demonstrated an important conservation in the DNA-binding preferences between *Sl*GBF3 and its Arabidopsis homolog, *At*GBF3. In Arabidopsis, *At*GBF3 is documented as a negative regulator of ABA response, with loss-of-function plants demonstrating an increased sensitivity to osmotic, drought and pathogen stress, and overexpressor plants showing reduced ABA sensitivity (Ramegowda *et al*., 2017; Dixit *et al*., 2019). Furthermore, Dixit et al. (2019) found that *At*GBF3 is essential for balancing drought tolerance and pathogen defense in the context of combined drought stress and *Pseudomonas syringae* infection.

Our findings suggest that *Sl*GBF3 has conserved its role in ABA-related responses, with a strong enrichment of genes involved in water deprivation, abiotic stimulus response and enriched response to ABA target genes. The GRNs showed that *Sl*GBF3 regulates important ABA-related genes, including *PP2C phosphatases*, *SnRK2 kinases*, and three ABF TFs (Fujii et al., 2009; Krukowski et al., 2023; Park et al., 2009). Notably, our study identified targets such as *Abscisic Acid Stress Ripening 1 (ASR1, Solyc04g071610)* and *dehydrin-like proteins (Solyc02g062390, Solyc02g084850*), which are indicators of ABA-mediated stress responses (Golan *et al*., 2014).

### Exploring tomato GRNs through the TomViz tools

The *Solanum lycopersicum* organ-level GRNs, available through the TomViz module on the PlantaeViz platform (Santiago et al., 2024), provide a robust and comprehensive resource for investigating TF-target interactions and organ-specific regulatory mechanisms. By integrating extensive datasets and emphasizing regulatory cascades, this tool surpasses traditional GCNs-based approaches, which are often limited to fruit tissue and a narrower gene set. The web application offers an accessible yet powerful platform for in-depth regulatory analysis, enabling researchers to explore tomato gene regulation across diverse developmental stages, environmental conditions, and genetic backgrounds. Its utility extends to various research contexts, facilitating novel discoveries in tomato biology and advancing functional genomics studies.

## Abbreviations

ABA: Abscisic acid
ABF: ABA response element binding factor
AUPR: Area under the precision-recall
AUROC: Area under the receiver operating characteristic
DAP-seq: DNA affinity purification sequencing
DEG: Differentially Expressed Gene
GENIE3: GEne Network Inference with Ensemble of Trees
GO: Gene Ontology
GCN: Gene coexpression network
GRN: Gene regulatory network
GSEA: Gene Set Enrichment Analysis
IVI: Integrated Value of Influence
OCS: Open Chromatin Sites
PP2C: Protein Phosphatase class 2 C.
PWM: Position Weight Matrix
RIN: RIPENING INHIBITOR
SnRK2: SNF1-related protein kinase 2.
TAGL1: TOMATO AGAMOUS-LIKE 1
TF: Transcription factor
TPM: Transcripts per Million

## Supplementary data

### Supplementary Figures

**Fig. S1.**
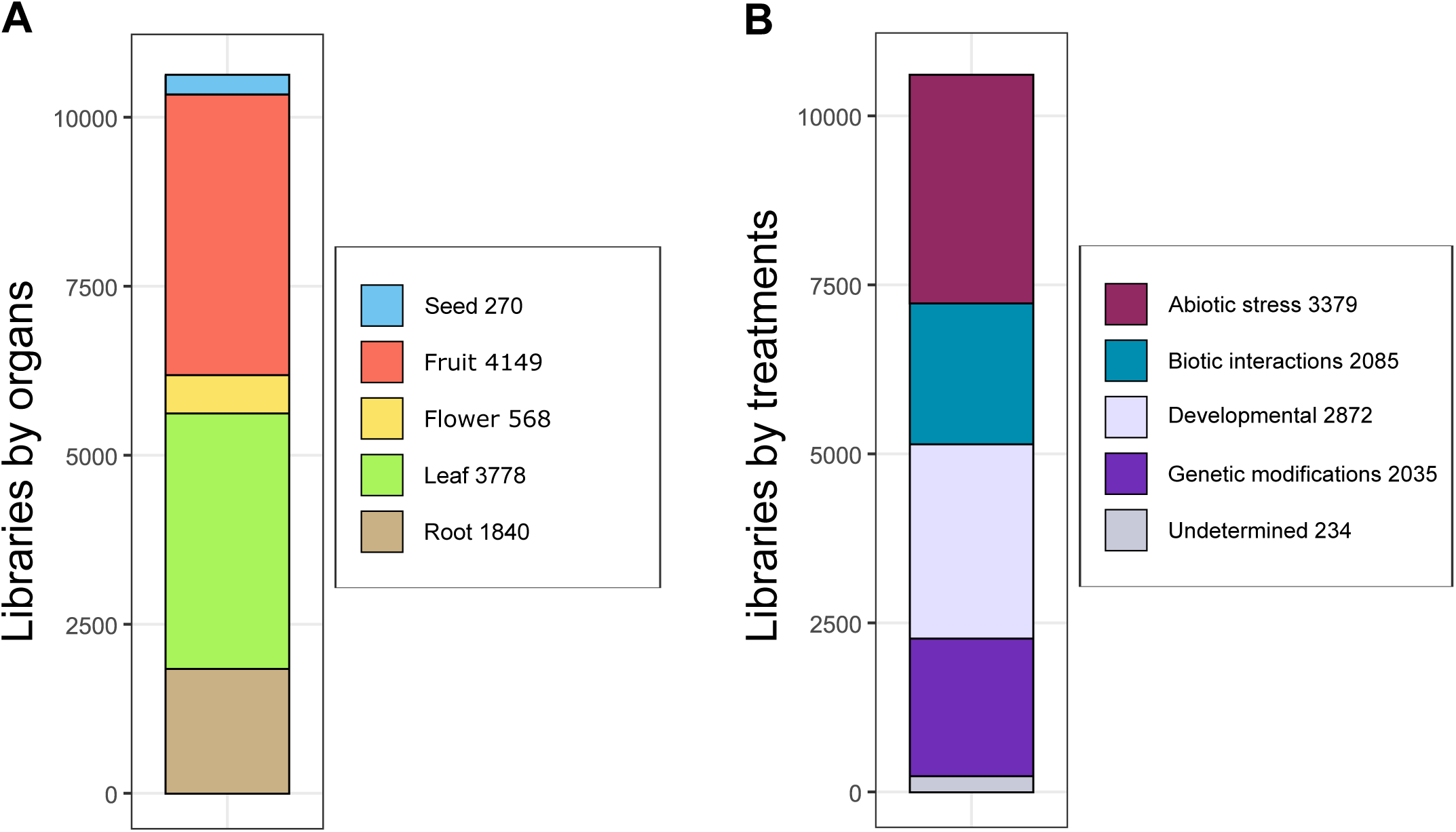
Distribution of tomato transcriptomic datasets across organs and experimental conditions. (A) Stacked bar plot showing the number of transcriptomic libraries classified by organ of origin (Seed, Fruit, Flower, Leaf, Root). (B) Stacked bar plot showing the distribution of transcriptomic libraries categorized by treatment or experimental condition (Abiotic stress, Biotic interactions, Developmental, Genetic modifications, Undetermined). The legend indicates the total number of studies per category.

**Fig. S2.**
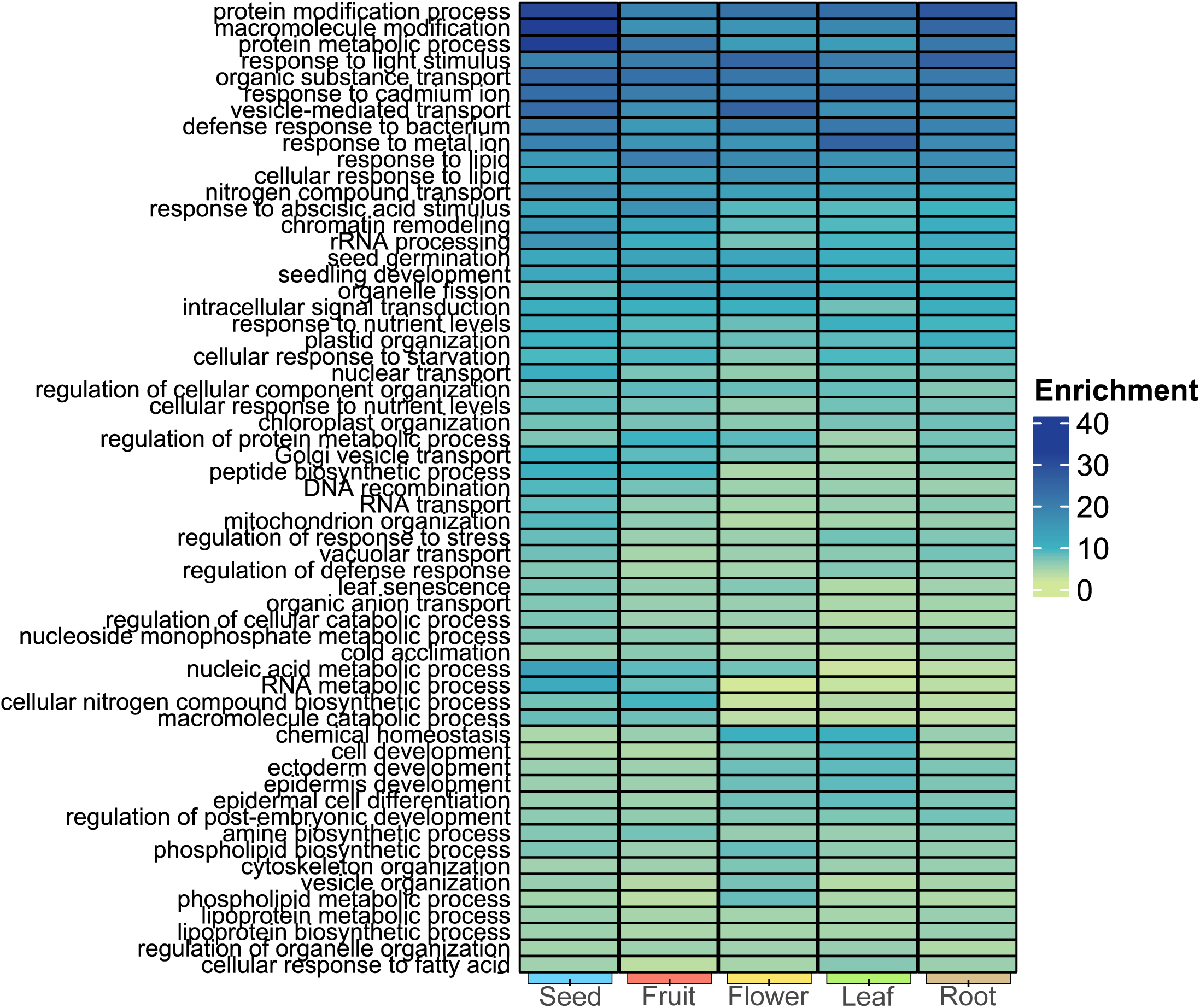
Heatmap of Gene Set Enrichment Analysis (GSEA) results for organ-specific gene expression (FDR-adjusted p-value < 0.05). Color intensity reflects the enrichment values.

**Fig. S3.**
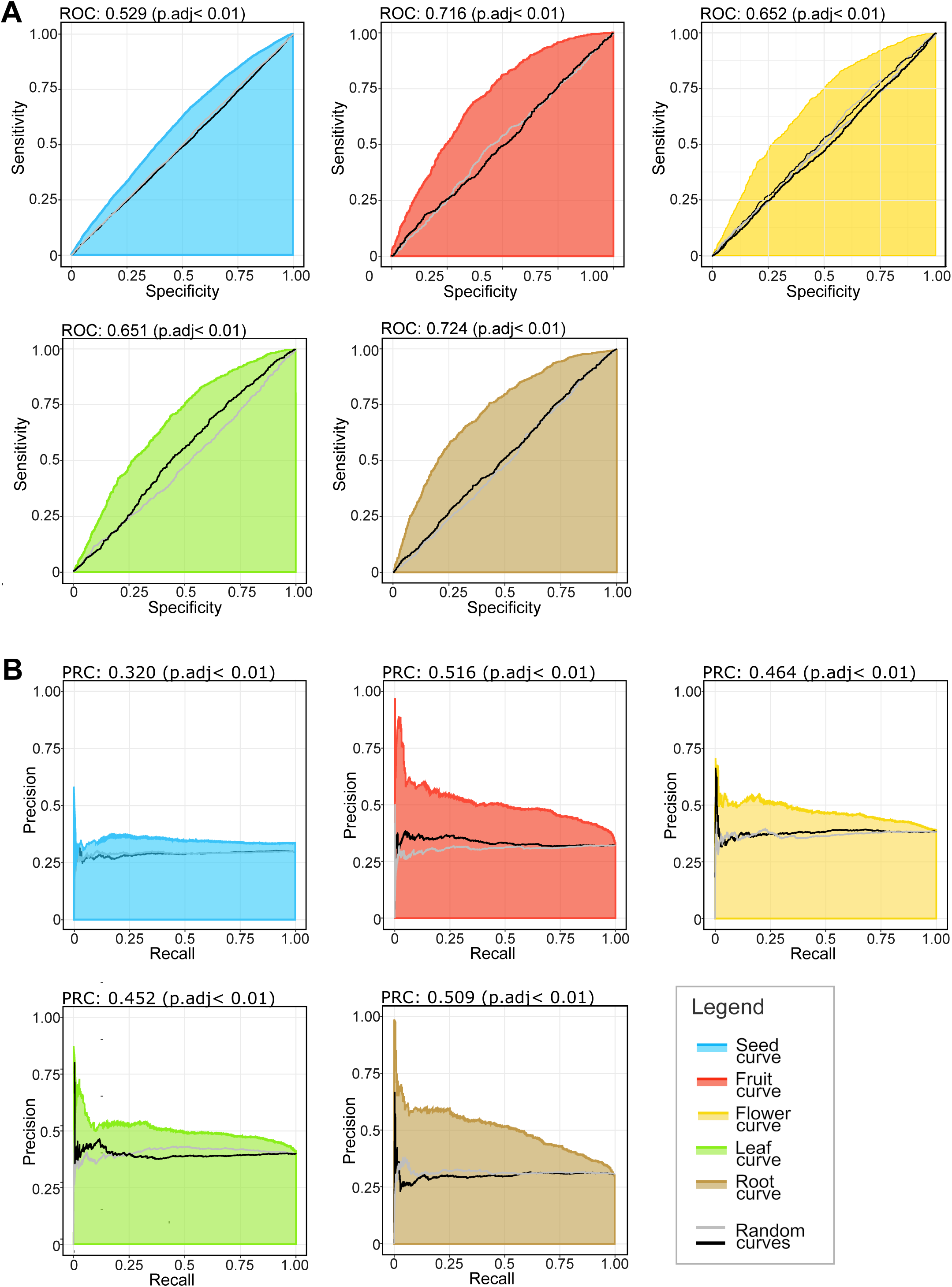
Accuracy analysis of organ-specific GRNs. Receiver Operating Characteristic (ROC) curves (A) and Precision-Recall (PR) curves (B) comparing organ-specific GRNs to ChIP-seq validation networks. The shaded areas indicate variability across multiple iterations. Black and grey lines represent the maximum and minimum quartiles of randomly generated TF-target pairings, respectively.

**Fig. S4.**
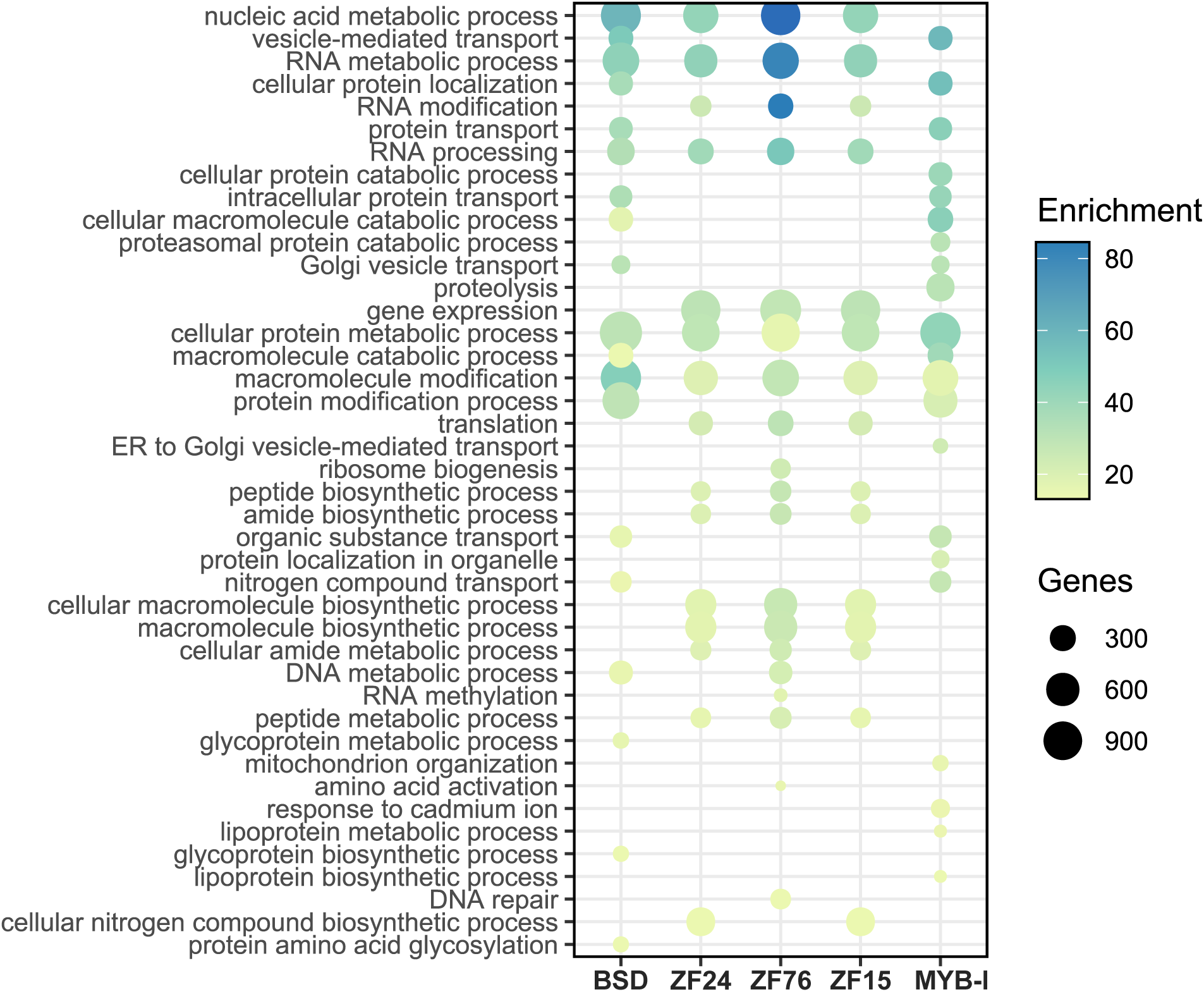
GSEA results (FDR-adjusted p-value < 0.05) for the five highest connected TFs shared across organ-level GRNs (conserved targets). Dot size represents gene number, while color intensity reflects enrichment values per TF.

**Fig. S5.**
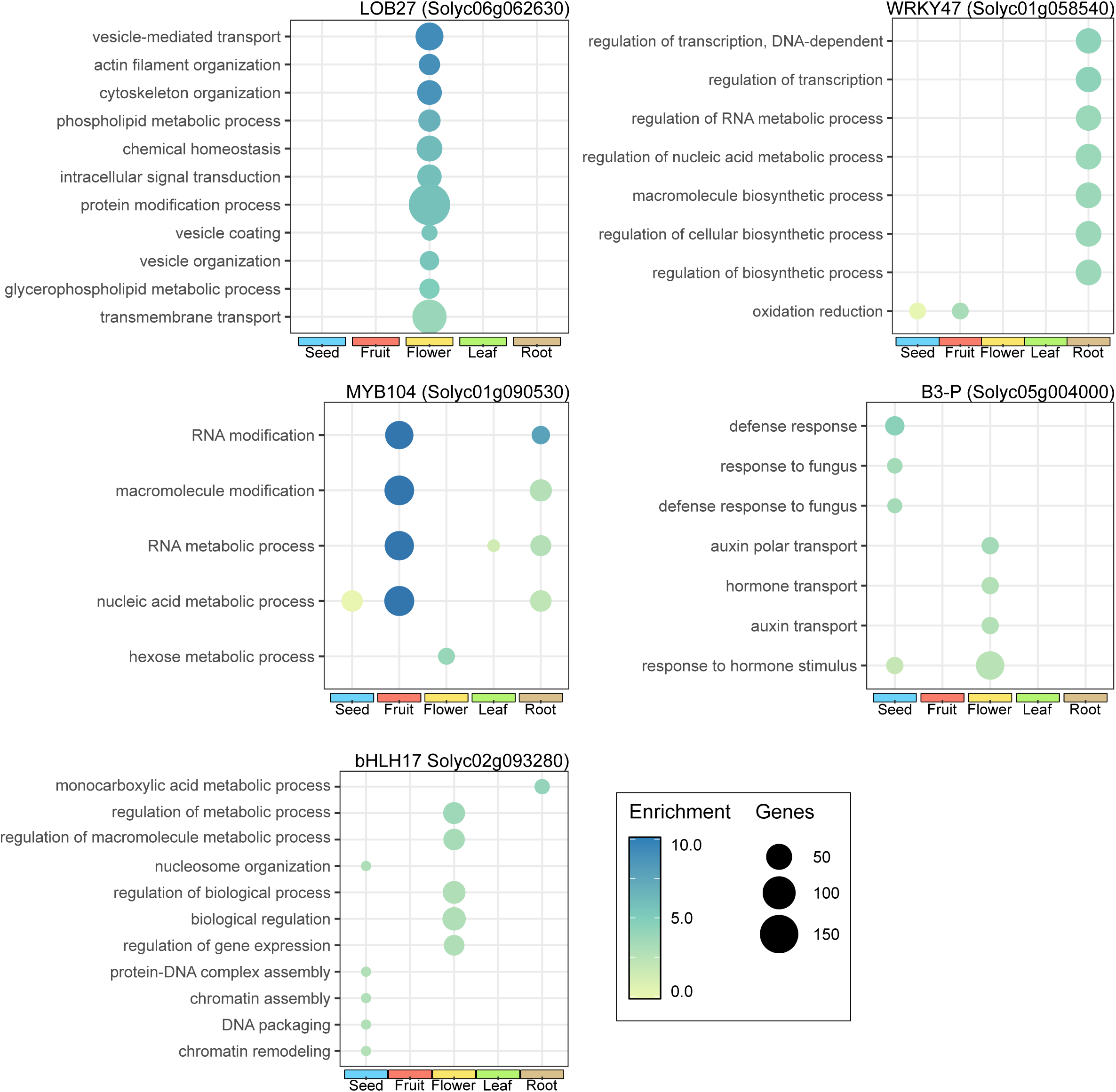
GSEA results (FDR-adjusted p-value < 0.05) for the five lowest connected TFs shared across organ-level GRNs (specific targets). Dot size represents gene number, while color intensity reflects enrichment values per TF.

**Fig. S6.**
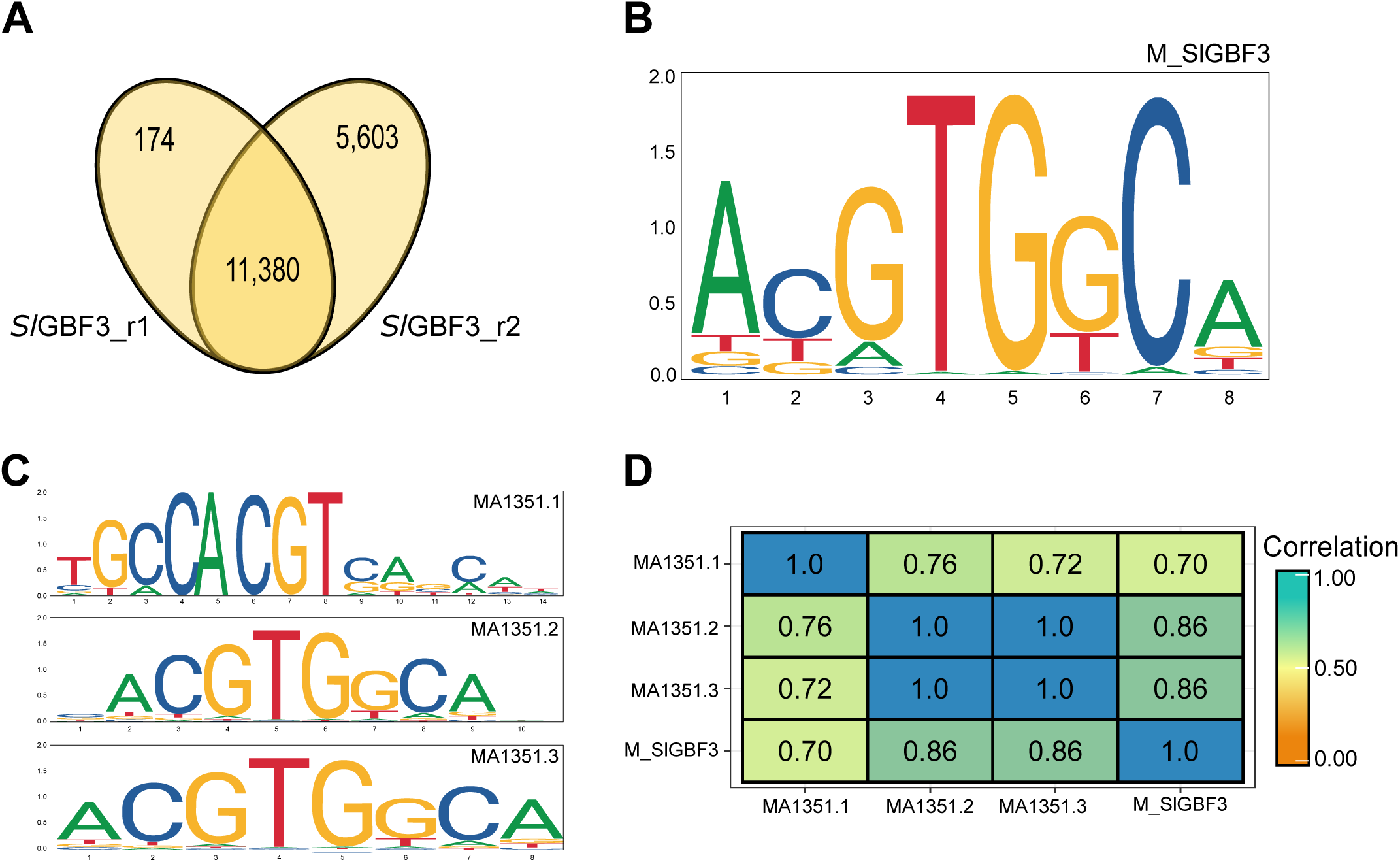
Genome-wide identification of *Sl*GBF3 binding targets using DAP-seq. (A) Venn diagram showing the overlap of DAP-seq peaks associated genes between two replicates (2). (B) *De novo* binding motif identified from *Sl*GBF3 DAP-seq peaks using the MEME suite (Machanick and Bailey, 2011). (C) Comparison of binding motifs from *At*GBF3 orthologs (MA1351.1, MA1351.2, MA1351.3). (D) Pearson correlation heatmap comparing the enrichment of *Sl*GBF3 binding motif (M_*SI*GBF3) with the *At*GBF3 orthologs.

**Fig. S7.**
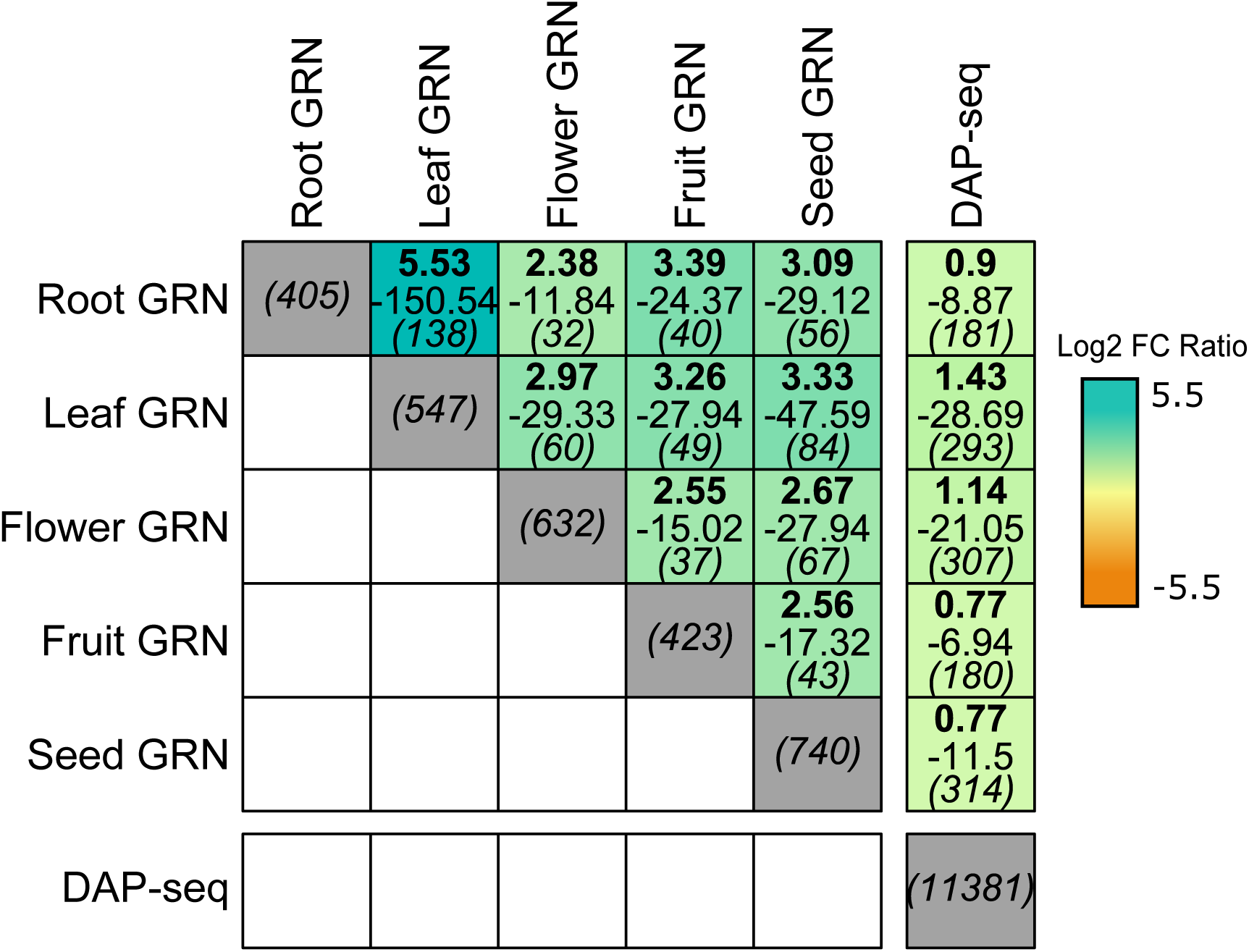
Enrichment analysis of *Sl*GBF3 targets in organ-specific GRNs compared to DAP-seq binding targets. Box heatmap display enrichment results from a Fisher’s exact test (log2 fold change, p-value, and intersection size).

**Fig. S8.**
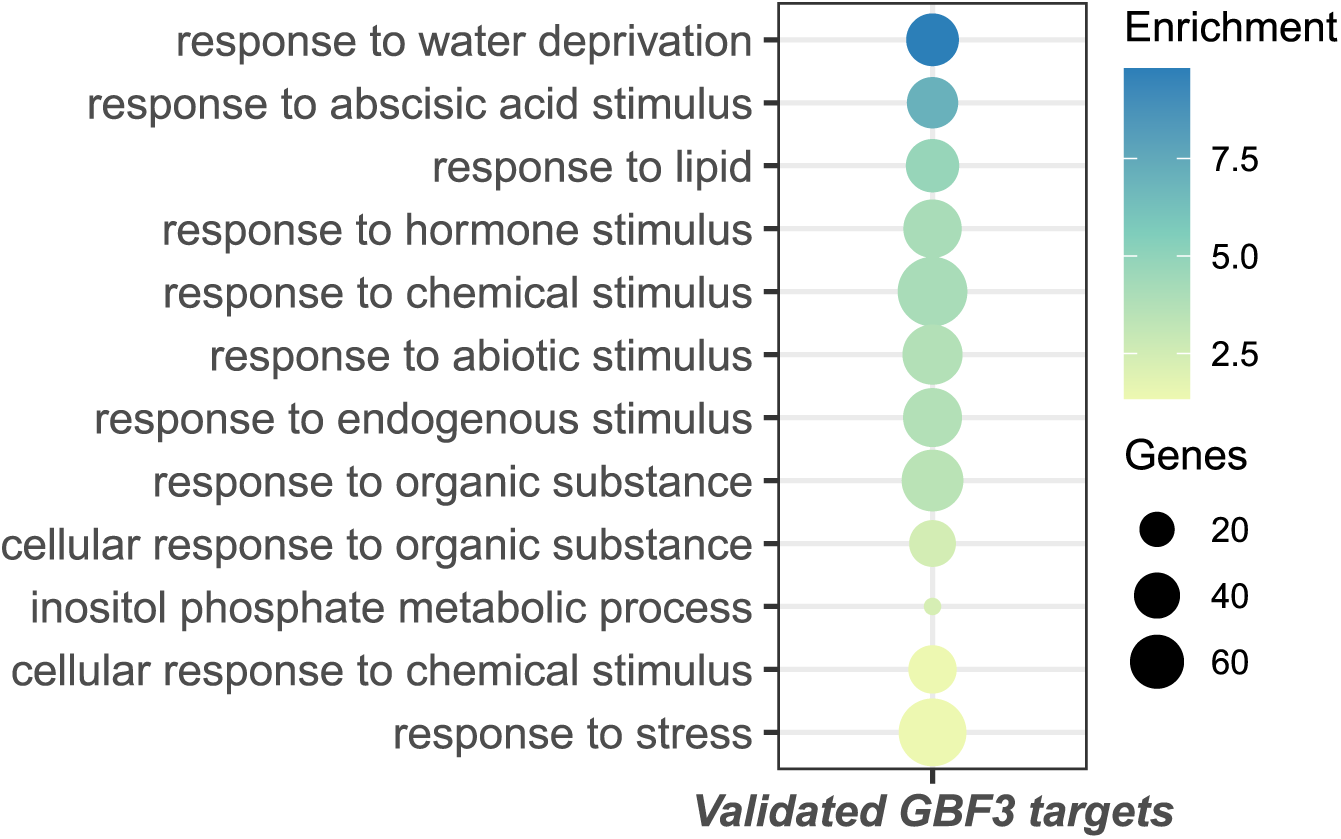
GSEA results (FDR-adjusted p-value < 0.05) of *Sl*GBF3 DAP-seq targets. Dot size represents gene number, while color intensity reflects enrichment values.

**Fig. S9.**
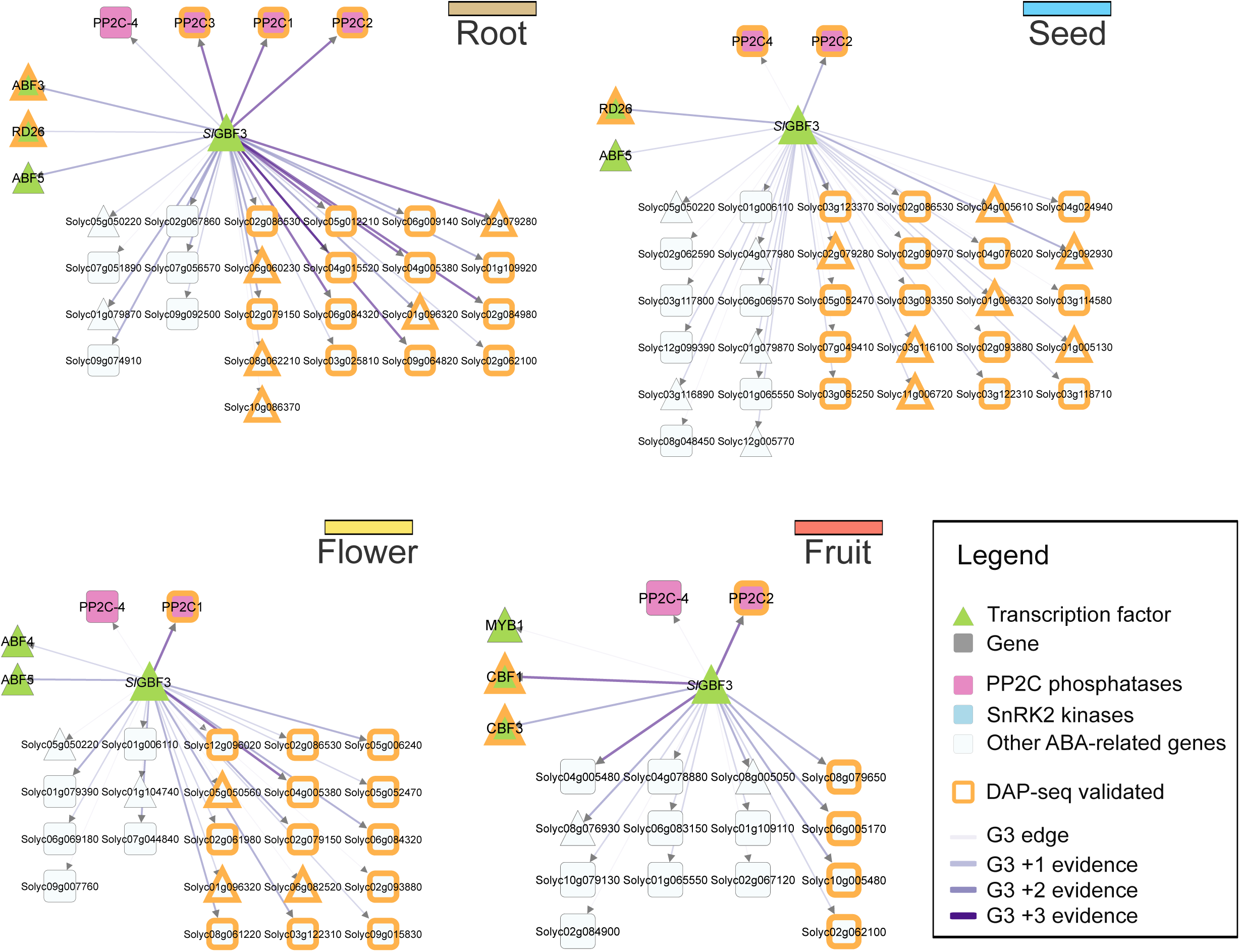
Network visualization of *SlGBF3*-regulated abscisic acid (ABA)-related genes within Root, Seed, Flower and Fruit GRNs. Triangles represent transcription factors, and squares represent genes. Node colors indicate function. Orange-bordered nodes highlight DAP-seq validated genes. Edge color shades represent accumulated regulatory evidence (G3: GENIE3).

### Supplementary Tables

**Supplementary Table S1.** Summary of Tomato Gene Annotations.

**Supplementary Table S2.** Gene Ontology (GO) annotations for *Solanum lycopersicum* (ITAG 4.0c).

**Supplementary Table S3.** Transcription factor (TF) evidence and position weight matrices (PWMs).

**Supplementary Table S4.** Transcriptomic libraries used for GRN construction.

**Supplementary Table S5.** Organ-level expression patterns.

**Supplementary Table S6.** Enriched GO terms shared across tomato organs.

**Supplementary Table S7.** Metadata of ChIP-seq libraries.

**Supplementary Table S8.** Enrichment analysis of GENIE3 predicted TF-target pairings.

**Supplementary Table S9.** Comparative enrichment analysis of GENIE3-predicted TF-target interactions and previously reported networks.

**Supplementary Table S10.** Metadata of Open Chromatin Sites (OCS) libraries.

**Supplementary Table S11**. TFs analysis reveals connectivity and target conservation distribution across tomato organs-level GRNs.

**Supplementary Table S12.** Ripening-related genes and their classification as RIN or TAGL1 targets.

**Supplementary Table S13.** Network analysis of key TFs in the fruit ripening network.

**Supplementary Table S14.** Genes annotated with the GO term "Response to ABA" or ABA-related genes.

**Supplementary Table S15.** ABA-related genes in tomato regulated by ABF TFs.

**Supplementary Table S16.** Leaf GRN targets of ABF3 and ABF5 regulating ABA-related and drought-responsive genes.

**Supplementary Table S17.** Network analysis of *Sl*GBF3 (Solyc01g095460) across all organ-specific GRNs.

**Supplementary Table S18.** DAP-seq analysis results for *Sl*GBF3.

## Acknowledgements

This research was supported by the computing infrastructure of the Center for Genomics and Bioinformatics, Universidad Mayor and the HPC cluster Garnatxa at the Institute for Integrative Systems Biology (I^2^SysBio). We thank the staff at SolGenomics (Dr. Surya Saha) for providing the iTAG4.2 beta annotation file.

## Author contributions

JDF, JMA, JTM and EAV: conceptualization; JDF, DN-P, AS: data curation; JDF, DN-P, AS, J Canan, SC-R, AC, TCM, LM and NRJ: methodology and investigation; JDF, DN-P, AS, TCM: formal analysis; EAV, JMA and JTM: funding acquisition; J Canales, JMA, JTM, and EAV: supervision; JDF, EAV: writing - original draft; JDF, J Canales, JMA, JTM and EAV writing - review and editing. All authors provided critical feedback and approved the final version of the manuscript.

## Data availability

The organ-level GRNs can be extracted and visualized through the GRN apps found within the TomViz module of the PlantaeViz platform (Santiago *et al*., 2024) available at https://plantaeviz.tomsbiolab.com/tomviz. The Supplementary Tables S4, S7, and S10 include the metadata of the public RNA-seq, ATAC-seq, DNASE-seq experiments, as well as the ChIP-seq libraries analyzed in this study. The DAP-seq data underlying this article is available at the NCBI Sequence Read Archive (SRA) database and can be accessed with accession number PRJNA1236412. The scripts used for analyzing RNA-seq, ATAC-seq, DNASE-seq, ChIP-seq, DAP-seq, the GENIE3 network generation pipeline as well of all the GRNs are available at: https://github.com/ibioChile/VidalLab/tree/master/Pipelines.

